# Distribution and dynamics of *Vibrio spp.* and pathogen sequences among microhabitats and monsoon seasons in Singapore’s coastal waters

**DOI:** 10.64898/2026.10.09.757811

**Authors:** Jeslyn S.T. Poo, Eric D. Hill, Patrick Martin, Janelle R. Thompson

## Abstract

Marine infections are rising in temperate regions due in part to a warming climate. However, comparatively little is known about drivers of marine pathogen dynamics in the tropics where temperatures are high year-round. *Vibrio* bacteria are ubiquitous in both temperate and tropical marine environments, and many are important pathogens of marine organisms and humans. Here, we sampled an equatorial monsoonal site, the Singapore Strait, over 18 months and applied amplicon sequencing (16s rRNA V6-8 region) to three particle size fractions, qPCR quantitation of *Vibrio* spp., and generalized linear modelling with environmental variables to understand the factors that structure the *Vibrio* assemblage. We found that Vibrios represented the highest proportion of pathogen-related DNA sequences, with an assemblage composition influenced by microhabitats (size fractions) and monsoon seasons. *Vibrio* sequences were dominated by ASVs with highest identity to *V. xuii*, *V. rotiferianus* and *V. caribbeanicus*, and including ASVs matching agents of human disease in Singapore – *V. parahaemolyticus*, *V. alginolyticus*, and *V. furnissii*. *Vibrio* qPCR abundance was lowest in the largest size fraction (10−150µm), following planktonic DNA trends, even though pathogen-related sequences, including *Vibrio*, were enriched in this fraction. Temperature was a significant predictor of overall *Vibrio* abundance, but its effect was no longer significant after adjustment for monsoons, suggesting that the temperature association may reflect broader seasonal dynamics. Instead, effects of chlorophyll, ammonia, nitrate and suspended solids were more consistent within seasons. Furthermore, free-living and particle-associated Vibrios responded differently to environmental changes, suggesting that partitioning into microspatial habitats may lead to differences in their environmental sensitivity.

## 1. Introduction

Marine pathogens, such as *Vibrio* species, are important in the context of climate change, as they are adapted to proliferate quickly in warm nutrient-rich environments, and already show rising infection numbers in recent decades (Baker-Austin et al. 2013). Vibrios typically constitute less than 1-3% of the oceanic bacterioplankton community (Rehnstam et al. 1993; Thompson et al. 2004) with numbers that vary significantly over regions and seasons. The genus contains more than 140 described species and four subspecies (Sampaio et al. 2022). Many have ecologically important roles in marine biogeochemical cycling (Grimes et al. 2009; Lin et al. 2018; Thompson and Polz 2006) and as symbionts and pathogens of diverse marine organisms. Twelve *Vibrio* species are known or emerging human pathogens (Baker-Austin et al. 2018; Osunla and Okoh 2017), and at least 25 are identified as causative agents of disease in wild and farmed marine fauna (de Souza Valente and Wan 2021; Mohamad et al. 2019; Sampaio et al. 2022). *Vibrio* abundance in seawater has therefore been proposed as a potential indicator of water quality and safety in coastal waters (Ng et al. 2018; Okpokwasili and Akujobi 1996; Purgar et al. 2023). Because of the direct association between warming temperatures and *Vibrio* abundance (Brumfield et al. 2025), the context of climate change highlights this group of bacteria as a growing global public and ecosystem health risk (Baker-Austin et al. 2017; Neoh et al. 2026).

*Vibrio* infections inflict significant economic and public health burden. For instance, estimated costs from *Vibrio*-induced acute hepatopancreatic necrosis in farmed shrimp exceed $43 billion USD since 2009 (Kumar et al. 2021; Shinn et al. 2018). In addition, *V. cholerae* is responsible for more than 100 000 global cholera-related deaths each year (Brumfield et al. 2025), and non-cholera Vibrios account for about 80 000 infections in the United States alone (Newton et al. 2012). In Singapore, *Vibrio* infections (other than Cholera) are not legally notifiable (Communicable Diseases Agency 2025), but reflect a higher per capita infection rate than the USA or Europe linked to the high relevance of seafood and seawater exposure as risk factors in Southeast Asia (Keng et al. 2025). Hospital data indicated that *Vibrio* infections in Singapore were mostly caused by *V. parahaemolyticus*, *V. fluvialis*, non-O1/non-O139 *V. cholerae*, *V. vulnificus* and *V. furnissii* (Neoh et al. 2026) with seasonal trends in relation to monsoons (Keng et al. 2025).

Several studies have examined Singapore’s coastal waters, observing the occurrence of Vibrios including *V. cholerae*, *V. parahaemolyticus* and *V. vulnificus* (Goh et al. 2017; Ivanov 2006; Ng et al. 2018). For instance, Goh et al. (2017) found overall higher prevalence of pathogenic *Vibrio* spp. in Singapore’s coastal waters relative to temperate studies and lowest frequency of toxigenic *V. parahaemolyticus* in more remote sites within the Singapore Strait (Goh et al. 2017). However, the factors that structure the *Vibrio* assemblage in the Singapore strait are currently unknown, i.e. whether it is influenced by the monsoon seasons, temperature, salinity or other physiochemical variables, and/or by microhabitat availability. Singapore’s coastal environment experiences variable water conditions influenced, for example, by high turbidity from coastal construction (Chou 1996; Guest et al. 2016; Low and Chou 1994), nutrient and pollutant loading from urban land-use (Browne et al. 2015; Sin et al. 2016), influx of colored dissolved organic matter (CDOM) from the regional peatlands (Martin et al. 2022), and episodic algal blooms (Hii et al. 2021; Leong et al. 2012; Trottet et al. 2018). The coastal environment is also subject to alternating wind-driven ocean currents due to the monsoon seasons, with currents flowing from the South China Sea during the Northeast (NE) monsoon (mid-November to March) and from the Malacca Strait during the Southwest (SW) monsoon (mid-May to September), separated by “inter-monsoon periods” (IM) of reduced flow through the Strait (Supplementary Fig. S1).

In this study we ask what factors structure the *Vibrio* assemblage in the Singapore Strait and whether the free-living and particle-associated populations follow similar or divergent trends. We have used 16s rRNA V6-8 gene amplicon sequencing, a group-specific *Vibrio* qPCR, measurements of seawater nutrients and physiochemistry, and generalized linear modelling, to quantify and model the abundance of *Vibrio* spp. in the Singapore Strait across three particle size fractions of seawater over 18 months. We hypothesized that bulk water physiochemical variables, microenvironments of suspended particles, and the monsoons seasons would influence the abundance and distribution of pathogens, allowing *Vibrio* abundance to be modelled and possibly predicted as a function of the environmental state. We also hypothesized that partitioning of *Vibrio* populations into microspatial habitats may lead to differences in the sensitivity of the populations to environmental drivers. Our results are, for the most part, consistent with both hypotheses. This study in a monsoon-driven environment complements previous studies in temperate (Eilers et al. 2000; Huang et al. 2024; Hunt et al. 2008; Szabo et al. 2013) and sub-tropical oceans (Liang et al. 2019; Wang et al. 2022) and extends the ability to model the response of Vibrios as an important marine pathogen group to a changing environment.

## 2. Materials and methods

### 2.1 Description of seawater sampling and metadata collection

Surface seawater was collected monthly from two sites in the Singapore Strait: St. John’s Island (1°12’59.0’’ N, 103°50’51.4’’ E) and Pulau Satumu (1°09’32.4’’ N, 103°44’23.3’’ E) over a period of 18 months (September 2021 to February 2023). Seawater was pre-filtered through a 150-µm plankton net and serially filtered through 10-µm, 1-µm and 0.22-µm polycarbonate membrane filters (Sigma-Aldrich, MO, USA; Sterlitech Corporation, WA, USA) to create three particle size fractions. Filtered samples were stored at –80°C before nucleic acid extraction.

At each sampling event, environmental parameters – temperature, salinity, dissolved nutrients (ammonia, NOx, nitrite, phosphate, silicate), dissolved organic carbon (DOC), dissolved inorganic carbon (DIC), total alkalinity, chlorophyll *a* and total suspended solids were measured according to Martin et al. (2022). The complete metadata across the 18-month time series is available at Poo et al. (2026).

### 2.2 DNA extraction and quantification

Genomic DNA was extracted from biomass collected on the membrane filters with the DNeasy PowerBiofilm Kit (Qiagen, Hilden, Germany). Samples were subjected to bead beating and chemical lysis and the extracted genomic DNA was cleaned with Genomic DNA Clean & Concentrator™ kit (Zymo Research, CA, USA). Purified DNA was quantified using Qubit™ 2.0 fluorometer and stored at –80°C until further molecular analyses.

### 2.3 Quantitative real-time PCR (qPCR) for quantifying Vibrio spp

A SYBR green qPCR assay targeting *Vibrio*-specific 16s rRNA gene (567F: 5’– GGCGTAAAGCGCATGCAGGT–3’; 680R: 5’–GAAATTCTACCCCCCTCTACAG–3’) (Thompson et al. 2004) was used to quantify *Vibrio* spp. abundance from each size fraction across the time series. The PCR cycling conditions were optimized based on Thompson et al. (2004) to achieve specific amplification of *Vibrio* and *Aliivibrio* spp. (collectively *Vibrio*) but not *Photobacterium* spp., which differed by a single nucleotide mismatch at each primer site. Annealing-elongation at 70°C achieved specific quantification of only *Vibrio* spp. with the TaqMan™ Fast Advanced Master Mix (Thermo Fisher Scientific, MA, USA). Assay specificity was validated by assessing amplification with a corresponding gene fragment from distantly related *Escherichia coli* and *Shewanella* sp. and the closely-related *Photobacterium* genus. Additionally, when challenged with a 50-fold excess of *Photobacterium* gene fragment, the assay yielded exclusively *Vibrio* product. The 567F/680R primer pair had an amplification efficiency of 86.7% under conditions optimized for *Vibrio* specificity and *Photobacterium* exclusion (Supplementary Fig. S2).

The absolute abundance of *Vibrio* spp. was presented as 16s rRNA gene copies of Vibrios per litre of seawater. Differences in the mean *Vibrio* 16s rRNA gene copies among the three size fractions were tested for statistical significance with the Kruskal-Wallis test, followed by Dunn’s test for post-hoc pairwise comparisons and multiple comparison correction. Results were considered statistically significant when *p* is less than 0.05.

### 2.4 Analysis of pathogen-containing genera sequences from Singapore time series 16s rRNA amplicon dataset

Amplicon sequencing was carried out using the primers B969F/BA1406 (B969F: 5’– ACGCGHNRAACCTTACC–3’, BA1406: 5’–ACGGGCRGTGWGTRCAA–3’ (Comeau et al. 2011)) targeting a 410–450 bp region of the 16s rRNA V6-8 gene. The amplicons were sequenced on the NovaSeq 6000 platform (PE250, Novogene) and sequence reads were processed with the DADA2 pipeline (Callahan et al. 2016) to generate amplicon sequence variants (ASVs), to which taxonomy was assigned by referencing the SILVA 16s rRNA database (v138). The dataset was then rarefied to 50 000 sequences per sample with total diversity discussed elsewhere (Poo et al. 2026). Here, we extract and analyze sequences classified to pathogen-pathogen-containing genera curated from four databases and additional resources: the US National Institute of Health Guidelines (National Institutes of Health 2024), the Pathosystems Resources Integration Center (Wattam et al. 2013) and an emerging infectious disease database (Jones et al. 2008), as well as from the aquaculture bacterial pathogen database (Lo et al. 2023), and from publications that identified putative pathogens responsible for coral diseases worldwide (Castañeda et al. 2018; Munn 2015; Sharma and Ravindran 2020; Vega Thurber et al. 2020). The final database used in this work consisted of 101 pathogen-containing genera and 329 listed pathogenic species (corresponding to known agents of human disease (including zoonoses) (43%) and animal disease (65%), noting that some species can infect both animals and humans). ASVs assigned to the genus *Vibrio* were further compared to the NCBI reference RNA collection (refseq_rna), excluding uncultured/environmental sample sequences, and the top search hit (highest BLAST score, lowest E value, highest percent nucleotide identity) is presented in Supplementary File 1, and when >99%, used as an ASV identifier.

### 2.5 Generalized linear models (GLMs) to relate Vibrio spp. abundance with environmental parameters and seasons

Two GLMs were constructed in R using the *glm* function to explore trends in *Vibrio* qPCR abundance in relation to the measured environmental parameters, i.e. Model 1: “Pooled,” where *Vibrio* abundance per sample was calculated as the sum of the measured concentrations from each size fraction and Model 2: “Size-fractionated”, where the size fractions are modelled separately. In an initial version of Models 1 and 2 (across-seasons), both models only accounted for the predictor effects of the measured environmental parameters and thus accounted for both across- and within-season variation. To test if predictor effects of the measured environmental parameters were significant within the monsoon seasons, both models were re-analyzed with ‘monsoons’ added as a categorical variable (within-seasons).

All models assumed a gamma distribution with a log-link function. Variables were natural-log transformed or mean-centered and those with highly correlations (NOx, phosphate, total alkalinity and DIC, rho ≥ 0.80) were removed to reduce multicollinearity, although we note some collinearity may remain, especially among monsoon-driven factors. For simplicity, no interactions were assumed for Model 1. In Model 2, interactions between variables and the categorical variable “size fraction” were determined. Non-significant interactions were excluded from the final version of Model 2.

The overall significance of terms in the models were assessed using a drop-one out analysis where the full model is compared to a model where each variable is removed one at a time using a Chi-squared test to assess change in model deviance and fit. The final model fit was evaluated by the Akaike Information Criterion (AIC) and the Bayesian Information Criterion (BIC), with lower values indicating a better fit (Supplementary Table S1). In addition, the Nagelkerke R-squared statistic (ranging from 0 to 1) was determined as a measure of how well the model explains the variance in the dependent variable. All contrast comparisons were adjusted for multiple comparisons using the Benjamini-Hochberg method. For comparison of trends, each ratio is estimated while holding all other variables constant.

### 2.6 GLMs to explore trends in the distribution of sequences matching human pathogenic Vibrios

To visualize temporal dynamics within the vibrioplankton, the relative abundance of *Vibrio* ASVs among total *Vibrio* sequences were transformed to environmental gene copy concentrations using the qPCR measurement of total *Vibrio* spp. obtained in this study. GLMs (Model 1 and Model 2 with and without control for seasons) were constructed for *Vibrio* ASVs matching human pathogens. Below sequencing limit of detection (LOD) data was accounted for by multiple imputation, and the final inference was then combined across imputations using Rubin-style pooling such that the reported coefficients, contrasts, confidence intervals, and *p* values reflect both ordinary sampling variation and the additional uncertainty introduced by imputation (van Buuren and Groothuis-Oudshoorn 2011) where Lambda is the fraction of total variance attributable to imputation uncertainty.

## 3. Results

### 3.1 Size-fraction distribution of sequences matching potential pathogens

Relative abundance of sequences from pathogen-containing genera was consistently higher in the largest particle size fraction (10–150 µm) relative to the free-living (0.22–1 µm; *p*=1.56e-6, 1.11e-5) and small-particle fraction (1–10 µm; *p*=3.23e-4, 1.50e-3) over 18 months at both St. John’s Island and Pulau Satumu (Fig. 1A). These sequences were also significantly more diverse in the 10–150 µm fraction than the 0.22–1 µm fraction (Fig. 1B, Shannon diversity: *p*=1.02e-4, 1.50e-3). Among sequences from pathogen-containing genera, *Vibrio* spp. comprised the majority (30.7 ± 15.6%, mean ± SD), followed by *Pseudoalteromonas* (14.2 ± 9.70%) and *Coxiella* (21.5 ± 16.3%; Fig. 1C). Occasional spikes of *Microbacterium* (0–46.9%) and *Halomonas* (0–28.3%) occurred in several samples (Fig. 1C).

**Fig. 1.**
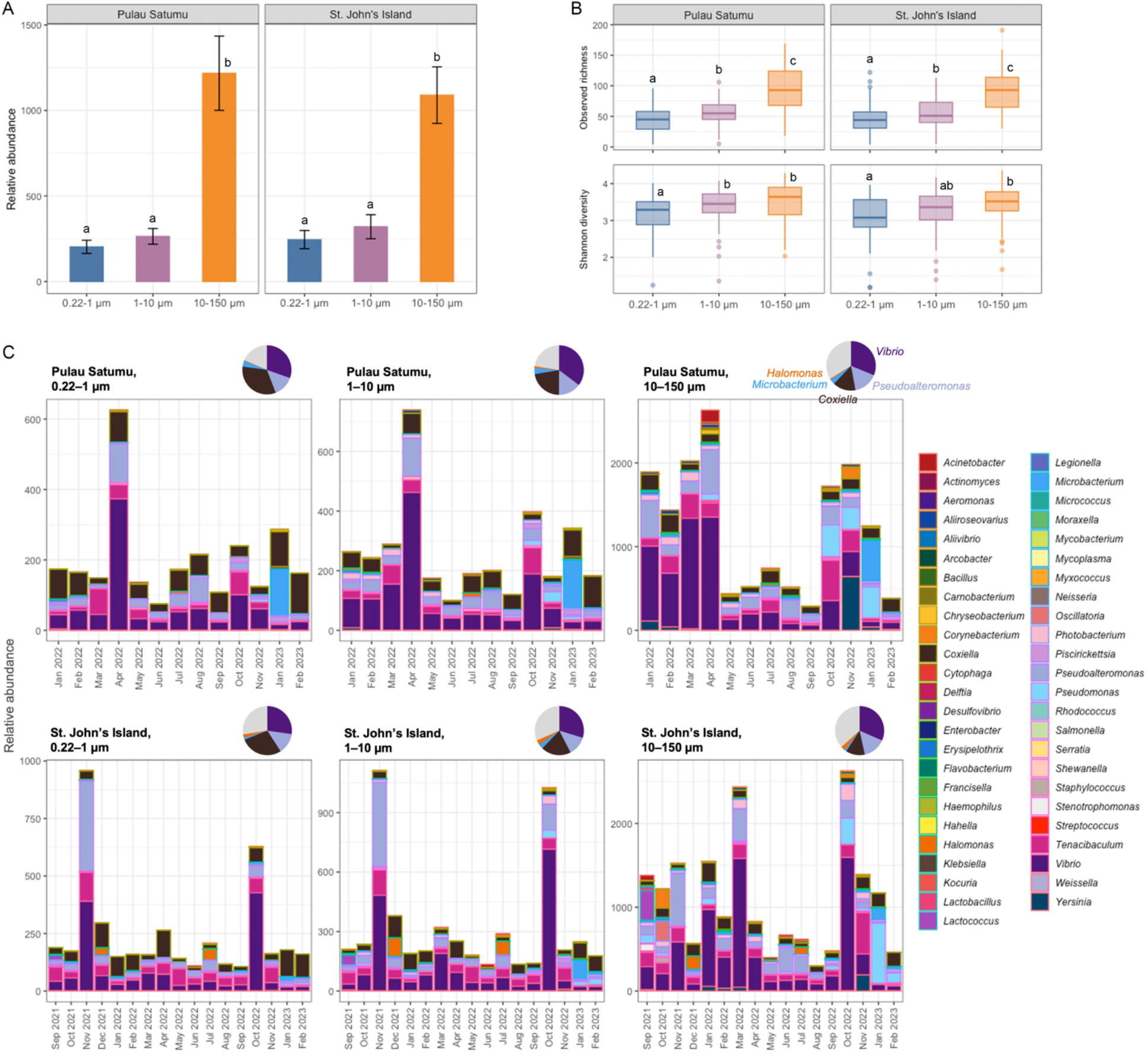
Distribution of ASVs from pathogen-containing genera among three size fractions in the Singapore Strait. (**A**) Relative abundance of sequences from pathogen-containing genera within the total bacterioplankton community (per 50 000 rarefied sequences) and (**B**) alpha diversity of these sequences in three size fractions at St. John’s Island and Pulau Satumu from the time series 16s rRNA amplicon dataset (Poo et al. 2026). Relative abundance and alpha diversity indices were compared for statistical difference among the three size fractions and groups that were significantly different are indicated by different letters (*p*<0.05). (**C**) Distribution (by relative abundance) of these genera among the three size fractions over the 18-month time series. Note the difference in scale of the vertical axes. The pie charts indicate average proportion of sequences from *Vibrio*, *Pseudoalteromonas*, *Coxiella*, *Microbacterium* and *Halomonas* over the time series (relative to all sequences from pathogen-containing genera).

Singapore Strait sequences contained 358 *Vibrio* ASVs mapping to 57 *Vibrio* species affiliations by BlastN to the Refseq database (Supplementary File 1). Although species-level assignments are tentative for amplicon sequences, the 404–405 bp V6-8 sequence region in *Vibrio* differentiates 99.5% of pairwise comparisons between reference *Vibrio* sequences in NCBI (Supplementary Fig. S3). The most abundant sequences on average were closely related (100% nucleotide identity) to *V. xuii* (ASV_214, *Vxu_214*), *V. rotiferianus* (ASV_237, *Vro_237*) and *V. caribbeanicus* (ASV_897, *Vca_897*). In the largest particle fraction (10–150 µm), the vibrioplankton was dominated by *Vxu_214* and *Vro_237*, together making up 34.4 ± 7.53% of the *Vibrio* assemblage at St. John’s Island and 32.5 ± 7.87% at Pulau Satumu (Supplementary Fig. S4). These two sequences had reduced relative abundances in the 0.22–1 µm fraction; instead *Vca_897* was dominant, constituting 15.1 ± 12.9% and 20.9 ± 18.2% at the two sites, respectively (Supplementary Fig. S4). Several of the most abundant *Vibrio* ASVs (Fig. 2) shared >99.5% nucleotide identity over the amplified V6-8 region with sequences from human pathogens *V. parahaemolyticus* (*Vpa*), *V. furnissii* (*Vfu*) and *V. alginolyticus* (*Val*) and marine and aquaculture pathogens such as *V. rotiferianus* (*Vro*) and *V. brasiliensis* (*Vbr*) (Supplementary File 1). Some of these pathogen-like ASVs were significantly enriched in the particle fractions, especially in the 10–150 µm fraction (*p*=0.00484−0.0446; Fig. 2).

**Fig. 2.**
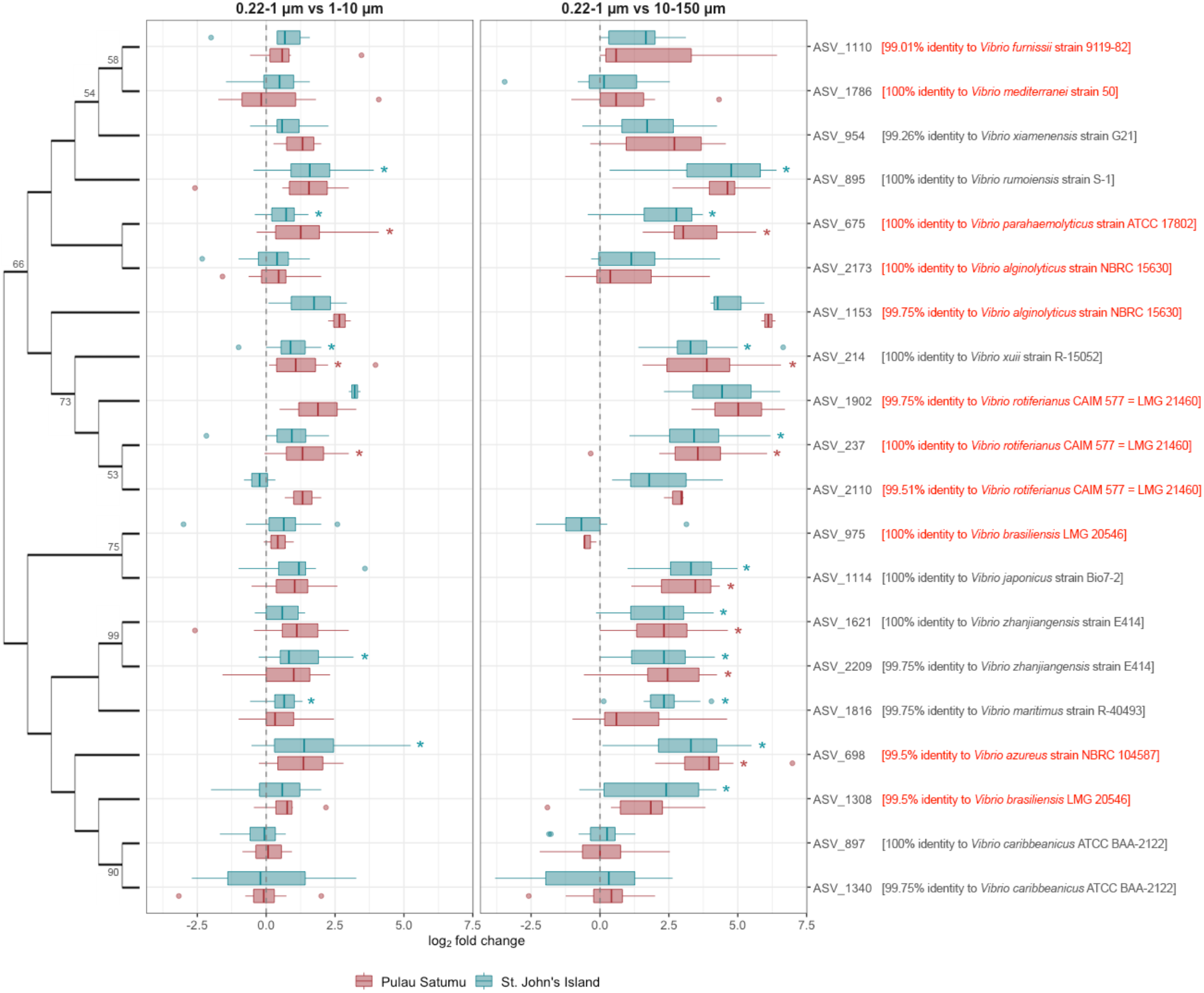
Relative proportion of the 20 most abundant *Vibrio* ASVs among the three size fractions. Enrichment was calculated as the log2-fold change of the relative abundance of sequences between two size fractions (see methods in Poo et al. (2026)). Positive values correspond to an enrichment in the particle-associated fraction (1–10 µm or 10–150 µm) while negative values correspond to enrichment in the free-living (0.22–1 µm) fraction. ASVs that were significantly enriched in a size fraction were indicated with asterisks (Wilcoxon signed-rank test with p-value adjustment using the Benjamini-Hochberg correction). These 20 ASVs were clustered based on their sequence identity over 404–405 bp using maximum likelihood approach (Tamura-Nei model, 1000 bootstrap replicates). Nodes with bootstrap support >50 are indicated. The top BLAST hit (against the NCBI Refseq database) of each ASV is also presented and those that were closely related to known *Vibrio* pathogens are indicated in red text.

### 3.2 Modelling total Vibrio spp. qPCR abundance in the Singapore Strait by GLM

Total *Vibrio* 16s rRNA gene copies reached 2.12 x 10^5^ to 4.87 x 10^6^ copies/L at St. John’s Island and 1.63 x 10^5^ to 5.68 x 10^6^ copies/L at Pulau Satumu (Fig. 3A), which were within the range of Vibrios quantified from seawater in Sweden (Eiler et al. 2006), USA (Thompson et al. 2004), Australia (Siboni et al. 2016) and China (Liang et al. 2019; Wang et al. 2020; Xu et al. 2021) with the same or similar assays. Across the seasons, total *Vibrio* abundance at both sites (Table 1) was significantly and positively associated with temperature (trend=1.70, BH-adjusted *p*=0.002), total suspended solids (trend=1.10, adjusted *p*=0.02) and chlorophyll *a* (trend=1.05, adjusted *p*=0.023), and negatively associated with nitrite (trend=0.884, adjusted *p*=0.023) and DOC (trend=0.975, adjusted *p*=0.01; Fig. 4A).

**Fig. 3.**
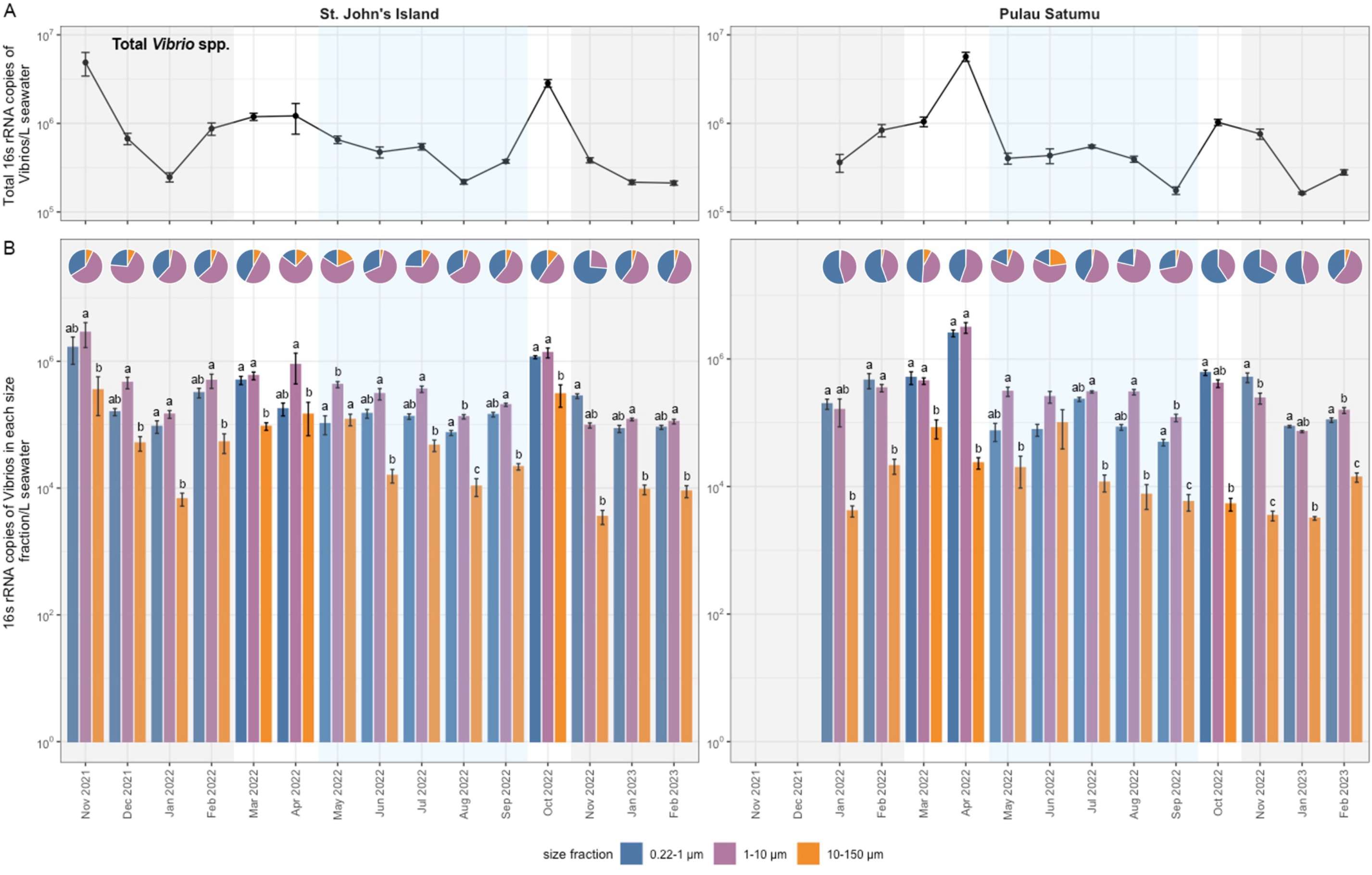
*Vibrio* spp. qPCR abundance quantified with the group-specific 567F/680R assay. Absolute abundance of (**A**) total *Vibrio* spp. (sum of *Vibrio* abundance from each size fraction) and (**B**) *Vibrio* spp. in the three size fractions at St. John’s Island and Pulau Satumu, presented as 16s rRNA copies of Vibrios per litre of seawater (*N*=5 per sampling time point and location). 16s rRNA copies of Vibrios were compared for statistical difference among the three size fractions at each month and groups with significantly different abundances are indicated by different letters (*p*<0.05). Monsoons and inter-monsoon periods are shaded with different background colors (NE monsoon: grey background, SW monsoon: blue background, IM-1 and IM-2: white background). For Pulau Satumu, the time series started in January 2022.

**Fig. 4.**
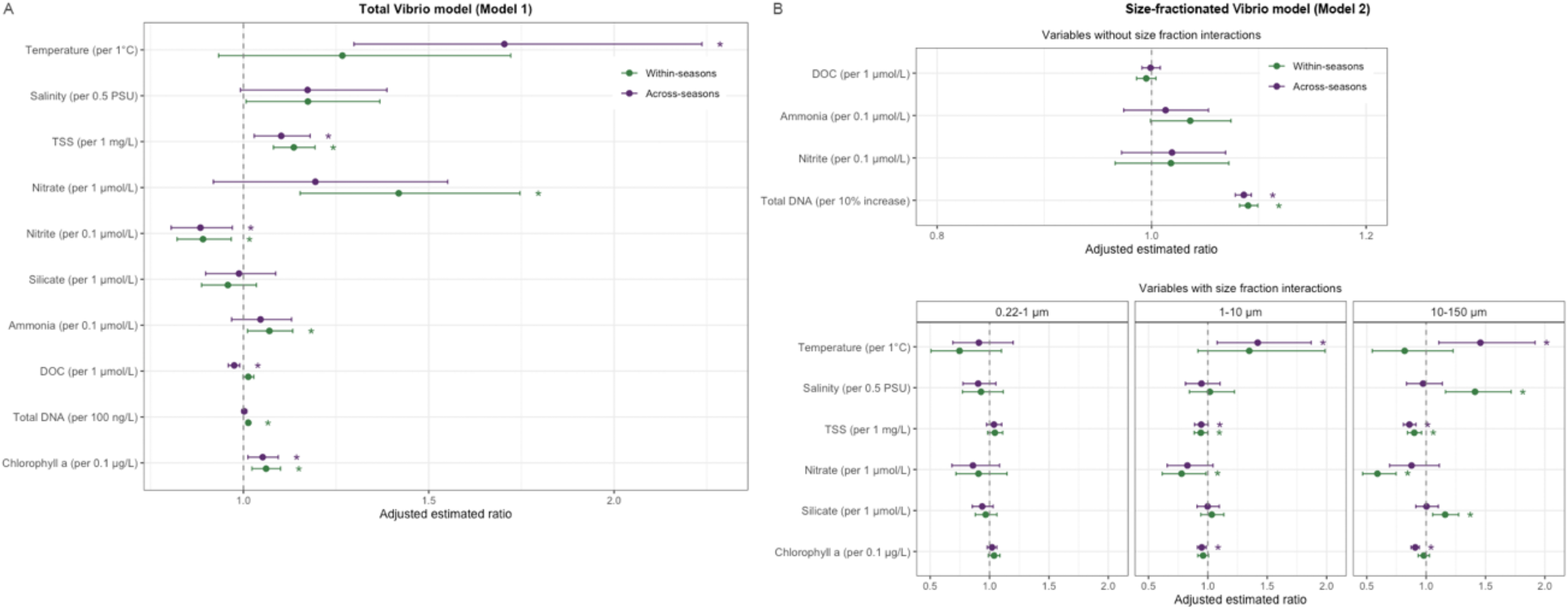
Comparison of the predictor effects of environmental parameters between the across-seasons model (without accounting for monsoons) and within-seasons model that included ‘monsoon seasons’ as a variable. Predictor effects were presented as adjusted estimated ratio with 95% confidence intervals for (**A**) total *Vibrio* spp. abundance (Model 1) and (**B**) *Vibrio* spp. abundance in each of the three size fractions (Model 2). Each trend is estimated while holding all other variables constant. All comparisons have been adjusted for multiple comparisons using the Benjamini-Hochberg method. Variables that were significant predictors (*p*<0.05) in each model are indicated with asterisks.

**Table 1.**
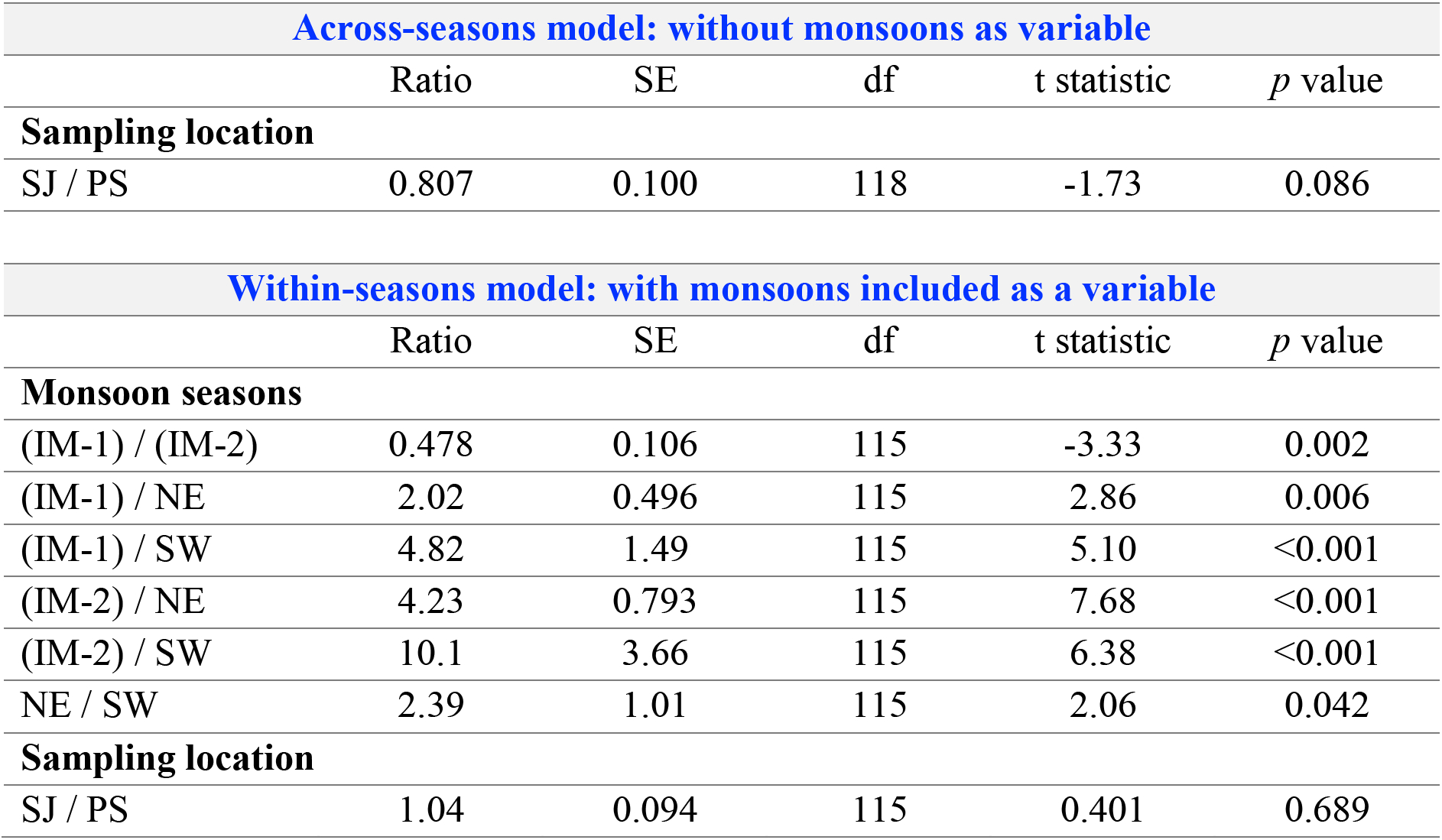
Estimates of categorical variables (monsoon seasons and sampling location) in the GLM analysis for total *Vibrio* abundance (sum of *Vibrio* abundance from each size fraction; Model 1). The model was first analyzed without accounting for seasonality (across-seasons) and then with monsoon seasons added as a separate variable (within-seasons). Each ratio is estimated while holding all other variables constant. All comparisons have been adjusted for multiple comparisons using the Benjamini-Hochberg method. NE: Northeast monsoon, IM-1: inter-monsoon 1, SW: Southwest monsoon, IM-2: inter-monsoon 2. SJ: St. John’s Island, PS: Pulau Satumu.

To test if seasonality influenced *Vibrio* distribution in this environment, the model was re-analyzed with ‘monsoon seasons’ included as a categorical variable, which improved the model fit (Supplementary Table S1) and emerged as a significant predictor of total *Vibrio* abundance (*p*<0.001, Supplementary Table S2). The inter-monsoon periods (IM-1 and IM-2) had 2- to 10-fold higher *Vibrio* concentrations than the NE and SW monsoons, and *Vibrio* abundance was lowest during the SW monsoon (Table 1). After accounting for monsoon seasonality, temperature and DOC were no longer significant predictors (temperature: adjusted *p*=0.142, DOC: adjusted *p*=0.081) whereas the effects of total suspended solids, nitrate, ammonia and total DNA became stronger (Fig. 4A, adjusted *p*<0.001, 0.004, 0.032, <0.001, respectively). In addition, chlorophyll remained highly significant within seasons (adjusted *p*=0.004), underscoring its importance as indicator of year-round primary productivity. We note that individual coefficients in the within-seasons model should be interpreted as conditional associations rather than independent effects due to residual collinearity among monsoon-associated parameters. The predictor effect of some monsoon-driven variables (e.g. temperature) could be encapsulated within the monsoon seasons, explaining why they appear to not exhibit significant effects on total *Vibrio* abundance in the model with seasonality (Fig. 4A).

### 3.3 Size-fraction trends in Vibrio spp. abundance

Most *Vibrio* DNA was found by qPCR in the 1–10 µm fraction at St. John’s Island and between the 1–10 µm and 0.22–1 µm fractions at Pulau Satumu (Fig. 3B). qPCR copies of *Vibrio* followed planktonic DNA trends, whose concentrations were lowest in the largest fraction (0.93–12.1% of pooled DNA per sample). Thus, despite enrichment of *Vibrio* ASVs in the 10– 150 µm fraction (Fig. 2), this fraction contained the lowest *Vibrio* spp. abundance by qPCR (Fig. 3B), making up <10% of the total *Vibrio* assemblage at St. John’s Island and <5% at Pulau Satumu. Significant positive correlations were noted between *Vibrio* qPCR abundance (16s rRNA gene copies/L) and *Vibrio* ASV relative abundance in the 0.22–1 µm and 1–10 µm fractions (r>0.8, *p*<0.001) with a weaker positive correlation for the 10–150 µm fraction (r=0.411, *p*=0.0299; Fig. 5) possibly due to higher variation in the more heterogeneous, particle-associated microenvironments.

**Fig. 5.**
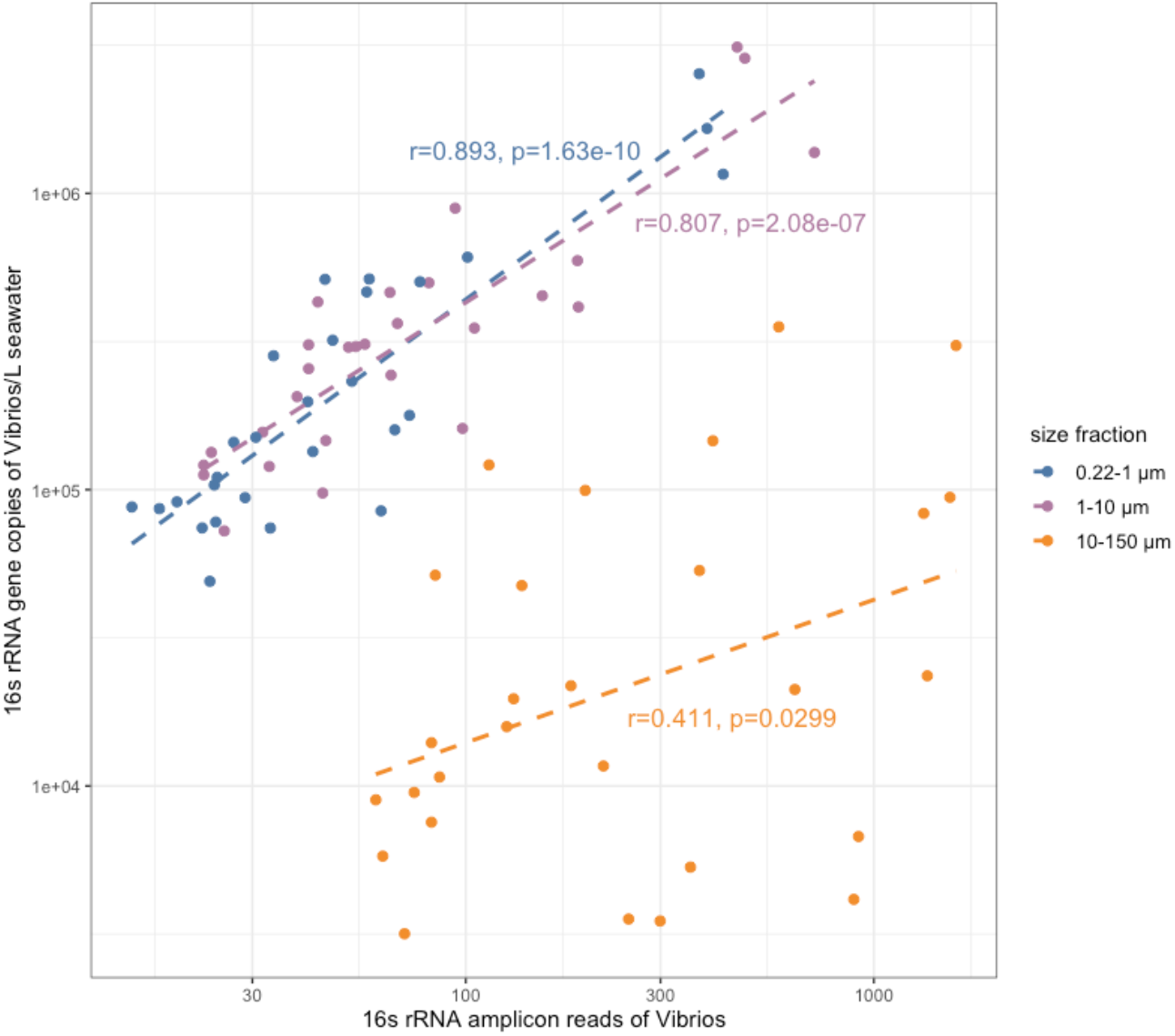
Correlation between the relative abundance of *Vibrio* 16s rRNA V6-8 sequences from the time series amplicon dataset and the qPCR abundance of *Vibrio* 16s rRNA gene copies. The Pearson’s correlation coefficient (r) and the significance of the correlation for each size fraction are indicated.

Significant interactions in the size-fraction explicit GLM (Supplementary Table S3) indicated size fraction-dependent responses of *Vibrio* spp. to some of the environmental variables considered in this study. In the across-seasons model, temperature, total suspended solids and chlorophyll *a* were significant predictors of *Vibrio* abundance in the intermediate and large particle size fractions (1–10 µm and 10–150 µm) but not in the “free living” 0.22–1 µm fraction (Fig. 4B). However, when monsoon seasonality was accounted for (within-seasons), seasonality emerged as an important predictor of *Vibrio* abundance among the size fractions. In fact, *Vibrio* abundance across the three size fractions varied significantly among the monsoon seasons (Table 2). During the NE monsoon and inter-monsoons, the highest *Vibrio* abundances were associated with the 0.22–1 µm and 1–10 µm fractions, reaching 17- to 38-fold higher than the 10–150 µm fraction (*p*<0.001, Table 2). However, during the SW monsoon, *Vibrio* abundance was more evenly distributed among the size fractions, accompanied by higher overall abundance in the largest size fraction (Table 2).

**Table 2.** Estimates of categorical variables and contrasts (monsoon seasons, sampling location and size fractions) in the GLM analysis for size-fractionated *Vibrio* abundance (Model 2). The model was first analyzed without accounting for seasonality (across-seasons) and then with monsoon seasons added as a separate variable (within-seasons). Each ratio is estimated while holding all other variables constant. All comparisons have been adjusted for multiple comparisons using the Benjamini-Hochberg method. NE: Northeast monsoon, IM-1: inter-monsoon 1, SW: Southwest monsoon, IM-2: inter-monsoon 2. SJ: St. John’s Island, PS: Pulau Satumu.

| Across-seasons model: without monsoons as a variable |  |  |  |  |  |  |
| --- | --- | --- | --- | --- | --- | --- |
| Contrast: size fraction | By sampling location | Ratio | SE | df | t statistic | p value |
| (0.22–1 $\mu\text{m}$) / (1–10 $\mu\text{m}$) | PS | 0.803 | 0.073 | 362 | -2.41 | 0.016 |
| (0.22–1 $\mu\text{m}$) / (10–150 $\mu\text{m}$) | PS | 11.5 | 1.06 | 362 | 26.6 | <0.001 |
| (1–10 $\mu\text{m}$) / (10–150 $\mu\text{m}$) | PS | 14.4 | 1.33 | 362 | 28.8 | <0.001 |
| (0.22–1 $\mu\text{m}$) / (1–10 $\mu\text{m}$) | SJ | 0.649 | 0.059 | 362 | -4.74 | <0.001 |
| (0.22–1 $\mu\text{m}$) / (10–150 $\mu\text{m}$) | SJ | 6.62 | 0.603 | 362 | 20.7 | <0.001 |
| (1–10 $\mu\text{m}$) / (10–150 $\mu\text{m}$) | SJ | 10.2 | 0.929 | 362 | 25.5 | <0.001 |
| Within-seasons model: with monsoons included as a variable |  |  |  |  |  |  |
| Contrast: size fraction | By monsoon seasons | Ratio | SE | df | t statistic | p value |
| (0.22–1 $\mu\text{m}$) / (1–10 $\mu\text{m}$) | IM-1 | 1.12 | 0.205 | 353 | 0.618 | 0.585 |
| (0.22–1 $\mu\text{m}$) / (10–150 $\mu\text{m}$) | IM-1 | 19.3 | 3.52 | 353 | 16.2 | <0.001 |
| (1–10 $\mu\text{m}$) / (10–150 $\mu\text{m}$) | IM-1 | 17.2 | 3.15 | 353 | 15.6 | <0.001 |
| (0.22–1 $\mu\text{m}$) / (1–10 $\mu\text{m}$) | IM-2 | 1.08 | 0.262 | 353 | 0.320 | 0.749 |
| (0.22–1 $\mu\text{m}$) / (10–150 $\mu\text{m}$) | IM-2 | 26.8 | 6.53 | 353 | 13.5 | <0.001 |
| (1–10 $\mu\text{m}$) / (10–150 $\mu\text{m}$) | IM-2 | 24.8 | 6.04 | 353 | 13.2 | <0.001 |
| (0.22–1 $\mu\text{m}$) / (1–10 $\mu\text{m}$) | NE | 0.652 | 0.196 | 353 | -1.42 | 0.186 |
| (0.22–1 $\mu\text{m}$) / (10–150 $\mu\text{m}$) | NE | 24.5 | 7.66 | 353 | 10.3 | <0.001 |
| (1–10 $\mu\text{m}$) / (10–150 $\mu\text{m}$) | NE | 37.6 | 11.5 | 353 | 11.9 | <0.001 |
| (0.22–1 $\mu\text{m}$) / (1–10 $\mu\text{m}$) | SW | 0.615 | 0.207 | 353 | -1.45 | 0.186 |
| (0.22–1 $\mu\text{m}$) / (10–150 $\mu\text{m}$) | SW | 1.94 | 0.663 | 353 | 1.94 | 0.080 |
| (1–10 $\mu\text{m}$) / (10–150 $\mu\text{m}$) | SW | 3.16 | 1.06 | 353 | 3.41 | 0.001 |
| **Contrast: monsoon seasons** | **By size fraction** | **Ratio** | **SE** | **df** | **t statistic** | **p value** |
| (IM-1) / (IM-2) | 0.22–1 $\mu\text{m}$ | 1.29 | 0.282 | 353 | 1.17 | 0.429 |
| (IM-1) / NE | 0.22–1 $\mu\text{m}$ | 1.61 | 0.405 | 353 | 1.89 | 0.153 |
| (IM-1) / SW | 0.22–1 $\mu\text{m}$ | 1.07 | 0.334 | 353 | 0.203 | 0.839 |
| (IM-2) / NE | 0.22–1 $\mu\text{m}$ | 1.25 | 0.260 | 353 | 1.06 | 0.429 |
| (IM-2) / SW | 0.22–1 $\mu\text{m}$ | 0.825 | 0.315 | 353 | -0.504 | 0.691 |
| NE / SW | 0.22–1 $\mu\text{m}$ | 0.662 | 0.297 | 353 | -0.920 | 0.461 |
| (IM-1) / (IM-2) | 1–10 $\mu\text{m}$ | 1.25 | 0.270 | 353 | 1.02 | 0.429 |
| (IM-1) / NE | 1–10 $\mu\text{m}$ | 0.936 | 0.240 | 353 | -0.256 | 0.839 |
| (IM-1) / SW | 1–10 $\mu\text{m}$ | 0.586 | 0.182 | 353 | -1.73 | 0.192 |
| (IM-2) / NE | 1–10 $\mu\text{m}$ | 0.751 | 0.162 | 353 | -1.33 | 0.372 |
| (IM-2) / SW | 1–10 $\mu\text{m}$ | 0.470 | 0.176 | 353 | -2.02 | 0.133 |
| NE / SW | 1–10 $\mu\text{m}$ | 0.625 | 0.279 | 353 | -1.05 | 0.429 |
| (IM-1) / (IM-2) | 10–150 $\mu\text{m}$ | 1.79 | 0.385 | 353 | 2.71 | 0.031 |
| (IM-1) / NE | 10–150 $\mu\text{m}$ | 2.05 | 0.559 | 353 | 2.62 | 0.033 |
| (IM-1) / SW | 10–150 $\mu\text{m}$ | 0.107 | 0.033 | 353 | -7.30 | <0.001 |
| (IM-2) / NE | 10–150 $\mu\text{m}$ | 1.14 | 0.263 | 353 | 0.576 | 0.678 |
| (IM-2) / SW | 10–150 $\mu\text{m}$ | 0.060 | 0.022 | 353 | -7.69 | <0.001 |
| NE / SW | 10–150 $\mu\text{m}$ | 0.052 | 0.024 | 353 | -6.43 | <0.001 |
| **Contrast: size fraction** | **By sampling location** | **Ratio** | **SE** | **df** | **t statistic** | **p value** |
| (0.22–1 $\mu\text{m}$) / (1–10 $\mu\text{m}$) | PS | 0.918 | 0.088 | 353 | -0.900 | 0.369 |
| (0.22–1 $\mu\text{m}$) / (10–150 $\mu\text{m}$) | PS | 15.7 | 1.52 | 353 | 28.5 | <0.001 |
| (1–10 $\mu\text{m}$) / (10–150 $\mu\text{m}$) | PS | 17.1 | 1.67 | 353 | 29.0 | <0.001 |
| $(0.22\text{--}1\ \mu\text{m}) / (1\text{--}10\ \mu\text{m})$ | SJ | 0.759 | 0.076 | 353 | -2.75 | 0.007 |
| $(0.22\text{--}1\ \mu\text{m}) / (10\text{--}150\ \mu\text{m})$ | SJ | 10.0 | 1.01 | 353 | 22.9 | <0.001 |
| $(1\text{--}10\ \mu\text{m}) / (10\text{--}150\ \mu\text{m})$ | SJ | 13.2 | 1.33 | 353 | 25.7 | <0.001 |

The within-seasons model showed that salinity, total suspended solids, nitrate and silicate also exhibited significant effects on *Vibrio* spp. abundance in the 10–150 µm fraction, but did not seem to influence the 0.22–1 µm fraction (Fig. 4B). Salinity had a significant positive effect (trend=1.41, adjusted *p*<0.001; Fig. 4B), suggesting that *Vibrio* spp. abundance in larger particles was elevated at higher salinity. Silicate also had a positive association with *Vibrio* spp. abundance in the 10–150 µm fraction (trend=1.16, adjusted *p*<0.001; Fig. 4B), potentially reflecting the role of silica-containing particles such as diatoms, radiolarians and smaller copepods on *Vibrio* abundance in the larger fraction. Total suspended solids and nitrate each had a significant negative association with *Vibrio* abundance in the 10–150 µm fraction (TSS trend=0.900, adjusted *p*<0.001; nitrate trend=0.590, adjusted *p*<0.001; Fig. 4B) despite positive effects on total *Vibrio* abundance (Fig. 4A). The significant interactions between size fraction and these variables suggest that *Vibrio* spp. abundance within different microhabitats in seawater respond differently to environmental and seasonal changes.

### 3.4 Dynamics of abundant Vibrio ASVs among the three size fractions and environmental parameters regulating their abundance

The abundance of individual *Vibrio* ASVs was estimated based on the qPCR measurement of *Vibrio* spp. transformed by their respective *Vibrio* sequence relative abundance (Supplementary Table S4) to explore whether the dynamics of individual *Vibrio* ASVs varied among the size fractions and monsoons. Despite higher qPCR abundance, *Vibrio* sequences in the 0.22–1 µm fraction were more variable where some ASVs were below detection on certain months, despite consistently appearing in the larger size fractions (Supplementary Fig. S5).

We found evidence of ecologically distinct populations within the vibrioplankton, defined as “groups of related strains that share a characteristic distribution among the sampled habitats” (Hunt et al. 2008; Preheim et al. 2011). We observed that similar *Vibrio* ASVs based on shared high 16s V6-8 identity to a reference sequence (e.g. *V. alginolyticus* (*Val*)) could vary in temporal dynamics over the study period (Supplementary Fig. S5). For instance, two *Val* ASVs (differing by 1 nt over the 405 bp sequence) alternated temporally with little overlap. *Val_1153* was detected sporadically (January – March 2022, October – November 2022, which corresponded to the inter-monsoons and the early or tail end of the NE monsoon) whereas *Val_2173* was detected throughout the time series but not before April 2022. Moreover, these two ASVs had distinct enrichment patterns, observed at both sampling sites (Fig. 2). This spatiotemporal pattern is consistent with the two *Val* ASVs reflecting distinct populations rather than different 16s rRNA operons.

On the other hand, in several cases closely related *Vibrio* ASVs had coherent environmental dynamics consistent with them being part of the same population, potentially due to intra-operon variation, noting that Vibrios may have up to 15 copies of the operon (Stoddard et al. 2015), sequences of which may vary up to 2% (Moreno et al. 2002) within a single strain. For example, three *V. rotiferianus* ASVs (ASV_237, ASV_1902 and ASV_2110) shared similar size fraction enrichment and temporal patterns (Fig. 2, Supplementary Fig. S5) and the same was observed for *V. zhanjiangensis* ASV_1621 and ASV_2209.

Among the top 20 most abundant *Vibrio* sequences, four were most closely related to human pathogens – ASV_675 (100% match to *V. parahaemolyticus* strain ATCC17802; *Vpa_675*), ASV_ 1110 (99.0% identity with *V. furnissii* strain 9119-82; *Vfu_1110*), ASV_ 2173 (100% match to *V. alginolyticus* strain NBRC 15630; *Val_2173*), and ASV_ 1153 (99.75% identity with *V. alginolyticus* strain NBRC 15630; *Val_1153*) (Supplementary Table S4). They were subjected separately to GLM analysis. The fraction of the total test variance attributable to imputation uncertainty from below-LOD data was 3% for *Vpa_675*, 24% for *Vfu_1110*, 23% for *Val_2173* and 39% for *Val_1153* (Supplementary Table S5). After accounting for imputation uncertainty, model fit was improved significantly by including monsoon seasons for *Vpa_675* (*p*=0.008), *Vfu_1110* (*p*<0.001) and *Val_1153* (*p*<0.001), but not for *Val_2173* (*p*=0.696). Trends below refer to the pooled fraction abundance estimates (Model 1) and the across-seasons GLM unless stated differently.

*V. parahaemolyticus*-like *Vpa_675* was observed to have the highest abundance during IM-1 (Fig. 6), and across seasons, seemed to be positively associated with salinity (*p*=0.006), ammonia (*p*=0.009) and chlorophyll (*p*=0.039; Supplementary Fig. S6A). All three variables appear to reflect seasonal rather than within-seasons variation as they become insignificant when monsoon season is controlled for (Supplementary Fig. S6A). In the size-fraction explicit model (Model 2), ammonia and DNA concentrations exerted significant positive effects on *Vpa_675* within seasons in all size fractions (*p*<0.001), with temperature (*p*=0.018) and total suspended solids (*p*<0.001) also contributing to within-seasons trends in the 1–10 µm fraction (Supplementary Fig. S6B).

**Fig. 6.**
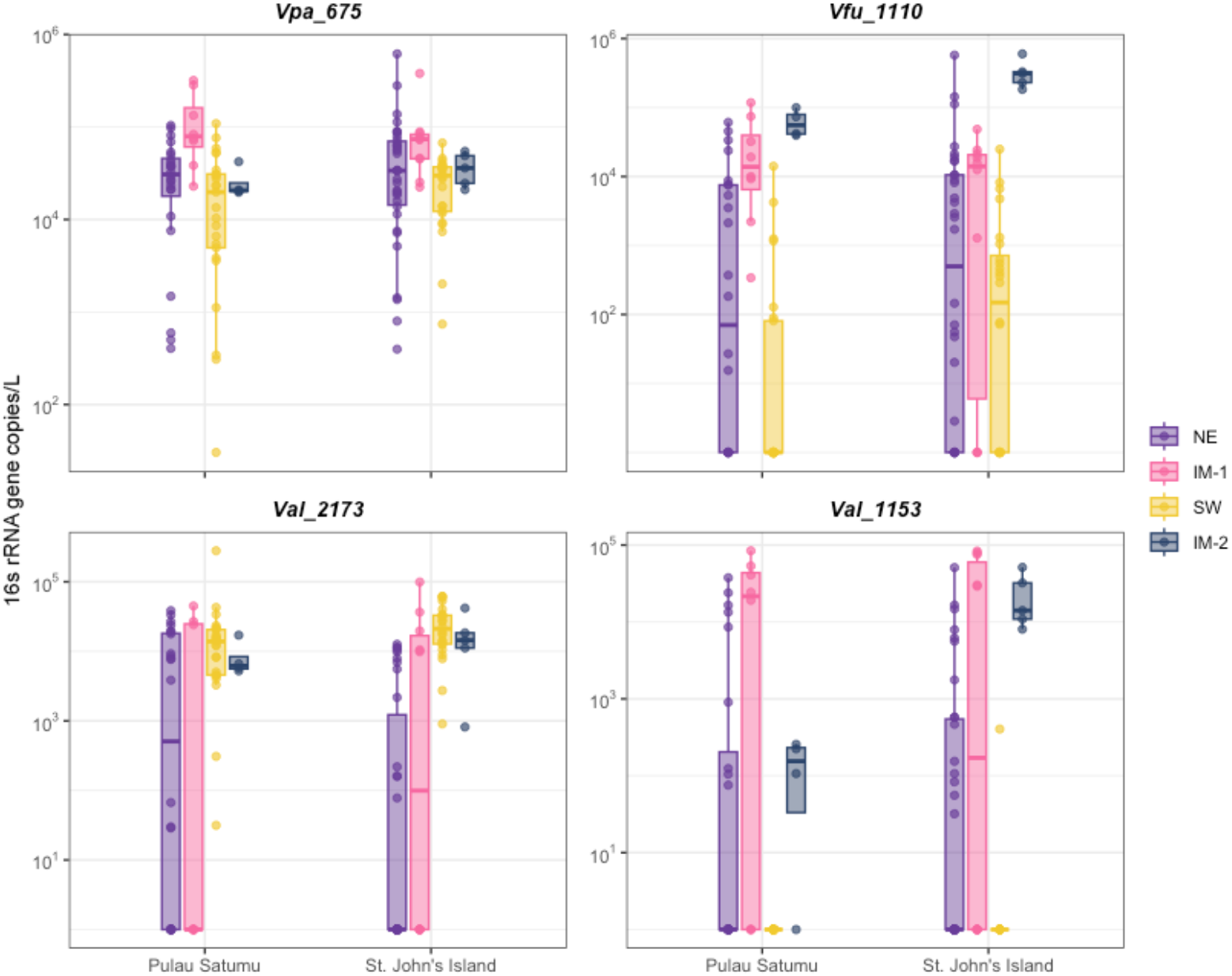
Abundance estimates of the four abundant *Vibrio* ASVs most closely related to human pathogens among the monsoon and inter-monsoon seasons. Abundance was calculated based on relative abundance from the time series 16s rRNA amplicon dataset and the qPCR *Vibrio* spp. abundance from each sample, plotted by monsoon periods – NE: Northeast monsoon, IM-1: inter-monsoon 1, SW: Southwest monsoon, IM-2: inter-monsoon 2.

The two *V. alginolyticus*-like ASVs (*Val_2173 and Val_1153*), which appeared to be ecologically differentiated based on particle size fraction association and temporal distribution (Fig. 2, 6), could not be fully compared in terms of their response to environmental variables due to a high proportion of imputed below-LOD data in the 0.22–1 µm fraction (Supplementary Table S5). Nevertheless, *Val_2173* could be shown to be positively influenced across- and within-seasons by total DNA (*p*=0.002 across seasons, *p*=0.003 within seasons) and nitrate (*p*<0.001), and negatively influenced by DOC (*p*<0.001 across seasons, *p*=0.007 within seasons; Supplementary Fig. S7A). Significantly positive influence of DOC and total DNA was observed in all size fractions, while the effect of nitrate was only significant in the largest size fraction (*p*=0.039; Supplementary Fig. S7B). By contrast, temperature emerged as a significant negative association for *Val_1153* in the within-seasons model (*p*=0.001) and this trend was reflected in all size fractions (Supplementary Fig. S8). In addition, within seasons, *Val_1153* was positively associated with salinity (*p*=0.025), total suspended solids (*p*=0.049) and chlorophyll (*p*=0.002; Supplementary Fig. S8A) with the salinity effect strongest in the largest size fraction (*p*<0.001; Supplementary Fig. S8B).

*V. furnissii*-like sequence *Vfu_1110* was at highest abundance during the inter-monsoons (Fig. 6). *Vfu_1110* abundance did not exhibit significant trends with the measured environmental variables likely due to high imputation associated variance (Supplementary Table S5). In the size fraction-explicit model (Model 2), *Vfu_1110* abundance in the largest size fraction was positively associated with salinity (*p*=0.016) and chlorophyll (*p*=0.003) and negatively associated with temperature within seasons (*p*<0.001; Supplementary Fig. S9), suggesting that the large particle-associated population may respond primarily to within-seasons variation rather than seasonal variation.

## 4. Discussion

### 4.1 Particles in the 10–150 µm fraction may constitute a source of pathogens in the marine environment

Despite observation that the largest (10–150 µm) size fraction had the lowest total DNA and *Vibrio* 16s rRNA gene copies in our study, we, and others (Bradshaw et al. 2016; Danovaro et al. 2009; He et al. 2022; Lyons et al. 2010), have observed an enrichment of pathogen-related sequences in association with particles, supporting the hypothesis that seawater particles are hotspots for microbial groups with pathogen traits and may be important for marine pathogen ecology. The mechanisms for the enrichment of pathogens in seawater particles are multi-fold. Particle enrichment may reflect pathogen origins in biomaterial shed by infected larger organisms, behavioural adaptations such as chemotaxis towards nutrient-rich microenvironments, and/or preferential growth at nutrient hotspots rich in labile organic compounds. The particle microenvironment may also serve to protect associated organisms against environmental stresses and predation (Russo et al. 2011). Moreover, some pathogenic microorganisms may have specific associations with eukaryotic plankton, such as the association between pathogenic *Vibrio* species and distinct diatom and copepod ASVs (Diner et al. 2021). Attachment to particulates (especially long-lived organic substrates such as marine snow or plankton) may enhance survival and transport of pathogens in marine environments (Craig et al. 2003; Haller et al. 2009), impacting their environmental persistence and virulence and has previously been identified as an important component of both the ecology and infection risk of certain pathogenic *Vibrio* species (Diner et al. 2021; Huq et al. 2005) in the context of past Cholera epidemics (Colwell 1996) and outbreaks of disease in mariculture such as *V. vulnificus* infection of the oyster *Crassostrea virginica* (Froelich et al. 2013). With a projected increase in seawater temperatures, which affect both *Vibrio* growth rates and particle coalescence (Danovaro et al. 2009), the possible expansion and persistence of pathogenic *Vibrio* populations housed within marine particulates is of particular concern when considering future projections of *Vibrio* infection risk.

The microbial community in the Singapore Strait is likely influenced by the advection of water masses through the Strait that varies with monsoon seasons, as well as by a combination of local factors. While the different water masses appear to carry distinct bacterioplankton communities (Chénard et al. 2019; Poo et al. 2026), local factors including sediment resuspension and shedding from benthic animals may occur seasonally or year-round. The balance of these sources to the structure of the *Vibrio* assemblage may differ across size fractions. For example, we have observed the largest size fraction associated ASVs exhibited more significant within-seasons trends with suspended solids, nutrients and total DNA (Fig. 4B, Supplementary Fig. S6-9), suggesting that particle-associated populations are more tightly coupled to localized environmental conditions that span seasons than their free-living counterparts.

### 4.2 Environmental factors shaping Vibrio spp. distribution in the Singapore Strait

Across studies (mostly in temperate regions), temperature and salinity have emerged as the strongest environmental correlates to *Vibrio* abundance (Takemura et al. 2014). In this study, across the monsoon seasons, total *Vibrio* abundance was significantly and positively associated with temperature over a narrow range of 4°C where SW monsoon and inter-monsoonal waters tend to be slightly warmer than NE monsoon waters (Martin et al. 2022; Poo et al. 2026; Tan et al. 2025). We suggest that changes in water masses associated with monsoonal reversal of ocean currents through the Strait may be more significant than the direct response of Vibrios to temperature changes. Indeed, if *Vibrio* spp. were responsive to temperature fluctuations in Singapore’s coastal waters, we would expect temperature to also emerge as a significant predictor of within-seasons variability. Nonetheless, temperature did emerge as a significant within-seasons predictor for *Val_1153*, which was also reflected across all fractions in the size fraction-explicit model, suggesting that certain populations closely related to human pathogens may be more responsive to local variations in temperature. Overall, finding of a diminished influence of temperature was consistent with other tropical studies (Asplund et al. 2011; Gregoracci et al. 2012; Sampaio et al. 2022; Takemura et al. 2014), indicating that other factors could be more important for regulating *Vibrio* abundance in warm tropical regions.

Seasonality is an important factor influencing *Vibrio* dynamics in this monsoon-driven tropical environment, as has been well-established for temperate regions. Here, total *Vibrio* abundance was significantly highest (2- to 10-fold) during the inter-monsoons and lowest during the SW monsoon (Table 1) and sequences most closely related to human pathogens *V. parahaemolyticus, V. alginolyticus* and *V. furnissii* followed these trends (Fig. 6). Finding of higher *Vibrio* abundance during the inter-monsoons was in contrast with studies that reported higher abundance during the wet monsoon season (Padovan et al. 2021; Simma et al. 2025; Vijayan and Lee 2014; Wong et al. 2019), which could be attributed to either higher temperatures (Simma et al. 2025) or higher precipitation and corresponding increase in land-based nutrient inputs (Vijayan and Lee 2014). The peak in *Vibrio* abundance during the inter-monsoons in Singapore’s coastal waters could instead be related to primary production, which is lowest during the early SW monsoon and peaks at the late SW and IM-2 (Zhang et al. 2026). We hypothesize that the increased primary production at the end of the SW monsoon (Zhang et al. 2026) releases bioavailable carbon and nutrients that may stimulate *Vibrio* growth and population expansion during IM-2.

Indeed, our GLM analysis revealed that chlorophyll *a* (related to primary production and phytoplankton biomass) was a strong positive predictor of total *Vibrio* abundance across and within seasons (Fig. 4A). Moreover, chlorophyll exhibited positive associations for *Vpa_675* in the free-living fraction and for *Val_1153* and *Vfu_1110* in larger size fractions, with some effects becoming stronger after controlling for seasons, thus indicating within-seasons drivers, especially for particle-associated populations. This finding was consistent with another coastal study in Southeast Asia (Wong et al. 2024), indicating that chlorophyll appears to reflect biologically meaningful, productivity-linked effects in tropical waters. On the other hand, surprisingly, DOC did not significantly influence total *Vibrio* abundance and in all three size fractions despite positive findings from temperate ecosystems (Eiler et al. 2007; Zhang et al. 2018), and *Val_2173* revealed a significantly negative trend that was robust to seasons. These results may partly be explained by the refractory nature of DOC in the Singapore Strait, where a large portion of terrigenous DOC originates from the surrounding peatlands in Sumatra (particularly during the SW monsoon) and is highly resistant to microbial degradation (Chu et al. 2025; Zhou et al. 2021).

Since Singapore’s waters are known for high turbidity, the relationship between *Vibrio* abundance and total suspended solids (TSS) was of particular interest. Suspended solids in Singapore’s seawater comprise mostly of fine-sized silica-rich clastic material (mean grain sizes 6.2−8.8 µm and 9.51−15.5 µm (Morgan et al. 2020)) and may also include organic particles (e.g. from plankton blooms or coral mucus). While measurements of TSS appeared to be positively related to across- and within-seasons variation of total *Vibrio*, this was qualitatively related to increases of free-living Vibrios as TSS was negatively associated with *Vibrio* abundance in the 1–10 µm and 10–150 µm fractions (Fig. 4). Explanation for the contrasting trends was unclear but could be related to the variable composition of seawater particles, which were not determined in this study. In particular, monsoonal winds, tidal currents and activities in Singapore’s seawater, such as coastal construction and ship traffic, may cause sediment resuspension introducing inorganic particles (Morgan et al. 2020) which may be less favorable to *Vibrio* attachment, than organic particles (Diner et al. 2021).

Inorganic nutrients that can be directly utilized for growth can also exert significant influence on *Vibrio* abundance (Kopprio et al. 2017; Thompson and Polz 2006). Singapore’s coastal waters are considered mesotrophic and have been shown to be nitrogen limited (relative to phosphorus) (Martin et al. 2022). We found that ammonia and nitrate levels had positive associations with total *Vibrio* abundance within seasons and ammonia levels showed a consistent and robust positive association with *Vpa_675* across all fractions. The nitrate effect was more variable across species/strains within the assemblage, with evidence of population-specific influence, for example, serving as a positive predictor for larger-particle associated *Val_2173*, both within- and across-seasons, due perhaps to roles of nitrate as a nutrient or anoxic electron acceptor.

Across studies, the patterns observed in the Singapore Strait samples both align with and diverge from established trends for human pathogenic Vibrios in tropical and temperate systems respectively. In temperate environments, *V. parahaemolyticus* and *V. alginolyticus* are typically strongly temperature-driven, with abundances peaking in warmer months and showing clear seasonal patterns tied to sea surface temperature and salinity gradients (Böer et al. 2013; Oberbeckmann et al. 2011; Takemura et al. 2014; Vezzulli et al. 2016). By contrast, the present results showed weak or absent temperature effects after controlling for monsoon seasonality, with temperature emerging only as a within-seasons predictor for *Val_1153*, suggesting that in tropical equatorial systems with minimal temperature variability, other drivers become more important. This aligns with tropical studies reporting reduced temperature sensitivity and greater influence of nutrients and productivity-related variables, such as chlorophyll and nitrogen species, on *Vibrio* dynamics (Asplund et al. 2011; Gregoracci et al. 2012).

Comparatively little is known about the environmental ecology of *V. furnissii*. Existing studies are largely limited to clinical reports (Ballal et al. 2017; Derber et al. 2011; Zhou et al. 2024), broad environmental surveys (Matté et al. 1994; Yibar et al. 2025) and some genomic studies (Wu et al. 2025), with few quantitative analyses of environmental drivers (Xu et al. 2020; Xu et al. 2021). Nevertheless, *V. furnissii* is recognized as an emerging pathogen associated with acute gastroenteritis (Ballal et al. 2017), typically acquired from contaminated water or seafood. The patterns observed here – particularly the association with chlorophyll and salinity in the largest size fraction and the emergence of within-seasons temperature effect provide new insights into the ecological controls on this understudied species.

While the analysis of *Vibrio* species diversity used in this study was limited by the choice of 16s rRNA as the marker gene, we demonstrated good discrimination of the 404−405 nt hypervariable V6-8 region of the 16S rRNA gene for *Vibrio* species in the Refseq database. We therefore tentatively interpret the presence of ASVs in the Singapore Strait that closely match pathogenic *Vibrio* species, including human pathogens, with a similar or possibly greater strength of evidence than what is provided by species-specific qPCR assays which generally target <100 nt of sequence information through binding of primers and probes. It is important to note that virulence and pathogenicity can be strain-specific and that 16s rRNA amplicon-based approaches cannot accurately distinguish virulent strains from less- or non-virulent ones. Nevertheless, 16S rRNA based approaches provide insights into the ecology of pathogen-related organisms, informing our understanding of how environmental changes may influence their distribution, dynamics, and ecology outside of the host niche including the environmental context for how virulence traits may be transmitted among closely related strains. In particular, traits such as cell size, motility and biofilm formation impact partitioning into microenvironments within the water column and are likely to be shared by closely related strains of variable virulence. While pathogenicity is determined based upon the presence and activity of specific virulence factors in the host-associated niche, and may be transferred horizontally among closely-related individuals (Eskova et al. 2022). Future work to identify pathogen traits and virulence genes among the size fractions, such as through metagenome sequencing, or genome sequencing of isolates, will be useful to better understand waterborne pathogen risk and to identify potential environmental drivers of selection for virulence genes outside of human or animal hosts.

## 5. Conclusion

In this study on *Vibrio* distribution and dynamics in the Singapore Strait, a key point of agreement with prior work is the importance of particle association. Previous studies in both temperate estuaries and tropical coastal waters have shown that *Vibrio* spp. are often enriched on organic particles, phytoplankton and zooplankton (Diner et al. 2021; Froelich et al. 2013; Main et al. 2015; Martinez-Urtaza et al. 2012), with chlorophyll and organic substrates serving as proxies for habitat and resource availability. The significant interactions between size fractions and some environmental parameters and the taxon-specific responses to nitrate, ammonia, chlorophyll and salinity are consistent with evidence that closely related *Vibrio* taxa occupy distinct ecological niches rather than responding uniformly to bulk environmental gradients. Overall, these results support a shift from temperature-dominated control in temperate regions to nutrient- and particle-driven structuring in tropical systems. Elevated seawater temperature has been shown to not only increase *Vibrio* abundance, but also to correlate with enhanced pathogenicity in some species where particle/plankton association alleviated the environmental constraints on the expression of virulence genes in some Vibrios (Billaud et al. 2022). In a particle-rich environment with a majority of Vibrios found within the 1–10 µm fraction and stable populations in the 10–150 µm fraction highlighting particles as important habitats, this could potentially reflect a pathway for enhanced pathogenicity, which has implications on marine ecosystem health. As Southeast Asian seas undergo projected climate shifts, our models suggest that monsoon-driven changes as well as local environmental changes could significantly impact *Vibrio* abundance in the Singapore Strait, including the abundance of notable human pathogens.

## Author contributions

**J.S.T. Poo**: conceptualization, data curation, formal analysis, investigation, methodology, software, visualization, writing – original draft, writing – review and editing. **E.D. Hill**: formal analysis, software, visualization, writing – review and editing. **P. Martin**: funding acquisition, investigation, resources, writing – review and editing. **J.R. Thompson**: conceptualization, formal analysis, funding acquisition, methodology, resources, supervision, writing – review and editing.

## Acknowledgements

This work was supported by the Singapore Ministry of Education and by the National Research Foundation, Singapore, through an RCE award to the Singapore Center for Environmental Life Sciences Engineering (SCELSE), start-up funding to J.R.T (Nanyang Technological University), the Marine Environment Sensing Network grant (NRF-NRI-2020-MESN) to P.M, and a research scholarship to J.P.S.T (Interdisciplinary Graduate Programme, Nanyang Technological University). The funders had no role in the study design, data collection or manuscript preparation. The physiochemical data collected as part of the Marine Environmental Sensing Network are available via https://ombak.mesn.sg.

## Data statement

Processed data and scripts will be made available upon request.

