## Supplementary for "Distribution and dynamics of *Vibrio spp.* and pathogen sequences among microhabitats and monsoon seasons in Singapore’s coastal waters"

Table S1. GLM fit statistics for Model 1 (“Pooled”: total *Vibrio* abundance summed across the three size fractions) and Model 2 (“Size-fractionated”: *Vibrio* abundance in each fraction). In both models, the model was first analyzed without accounting for seasonality (across-seasons) and then with monsoon seasons added as a separate variable (within-seasons).

| Statistic | Across-seasons model | Within-seasons model |
|---|---|---|
| **Model 1 (“Pooled”)** Observations: 130 | | |
| AIC | 3662.810 | 3576.409 |
| BIC | 3700.088 | 3622.289 |
| Nagelkerke R-squared | 0.721 | 0.863 |
| **Model 2 (“Size-fractionated”)** Observations: 420 | | |
| AIC | 9497.403 | 9440.975 |
| BIC | 9612.421 | 9591.689 |
| Nagelkerke R-squared | 0.927 | 0.939 |

Table S2. Overall significance of terms in the GLM analysis for total *Vibrio* abundance (Model 1). The overall significance of terms were assessed using a drop-out analysis (see methods). The model was first analyzed without accounting for seasonality (across-seasons) and then with monsoon seasons added as a separate variable (within-seasons).

| | df | Deviance | AIC | Scaled deviance | *p* (>Chi) |
|---|---|---|---|---|---|
| **Across-seasons model: without monsoons as a variable** | | | | | |
| Total Model | - | 39.0 | 3662.810 | - | - |
| Temperature | 1 | 43.7 | 3675.240 | 14.4 | <0.001 |
| (Salinity/0.5) | 1 | 40.2 | 3664.338 | 3.53 | 0.060 |
| Sampling location | 1 | 40.0 | 3663.915 | 3.11 | 0.078 |
| Total suspended solids | 1 | 41.6 | 3668.645 | 7.84 | 0.005 |
| Nitrate | 1 | 39.6 | 3662.495 | 1.69 | 0.914 |
| (Nitrite/0.1) | 1 | 41.3 | 3667.638 | 6.83 | 0.009 |
| Silicate | 1 | 39.1 | 3660.871 | 0.061 | 0.805 |
| (Ammonia/0.1) | 1 | 39.4 | 3661.934 | 1.13 | 0.289 |
| DOC | 1 | 42.0 | 3669.988 | 9.18 | 0.002 |
| (Total DNA/100) | 1 | 39.2 | 3661.321 | 0.511 | 0.475 |
| (Chlorophyll *a*/0.1) | 1 | 40.5 | 3665.370 | 4.56 | 0.033 |
| **Within-seasons model: with monsoons included as a variable** | | | | | |
| Total Model | - | 19.7 | 3576.409 | - | - |

| | | | | | |
|---|---|---|---|---|---|
| Temperature | 1 | 20.0 | 3576.705 | 2.30 | 0.130 |
| (Salinity/0.5) | 1 | 20.3 | 3578.629 | 4.22 | 0.040 |
| Sampling location | 1 | 19.7 | 3574.573 | 0.164 | 0.685 |
| Monsoon seasons | 3 | 39.0 | 3693.728 | 123 | <0.001 |
| Total suspended solids | 1 | 23.6 | 3599.772 | 25.4 | <0.001 |
| Nitrate | 1 | 21.4 | 3585.747 | 11.3 | <0.001 |
| (Nitrite/0.1) | 1 | 21.0 | 3582.647 | 8.24 | 0.004 |
| Silicate | 1 | 19.9 | 3575.644 | 1.24 | 0.266 |
| (Ammonia/0.1) | 1 | 20.5 | 3579.474 | 5.07 | 0.024 |
| DOC | 1 | 20.2 | 3577.698 | 3.29 | 0.070 |
| (Total DNA/100) | 1 | 24.4 | 3604.537 | 30.1 | <0.001 |
| (Chlorophyll *a*/0.1) | 1 | 21.2 | 3584.293 | 9.88 | 0.002 |

Table S3. Overall significance of terms in the GLM analysis for size-fractionated *Vibrio* abundance (Model 2). The overall significance of terms were assessed using a drop-out analysis (see methods). The model was first analyzed without accounting for seasonality (across-seasons) and then with monsoon seasons added as a separate variable (within-seasons).

| | df | Deviance | AIC | Scaled deviance | *p* (>Chi) |
|---|---|---|---|---|---|
| **Across-seasons model: without monsoons as a variable** | | | | | |
| Total Model | - | 85.1 | 9497.403 | - | - |
| DOC | 1 | 85.1 | 9495.421 | 0.018 | 0.894 |
| (Ammonia/0.1) | 1 | 85.2 | 9495.822 | 0.419 | 0.517 |
| (Nitrite/0.1) | 1 | 85.2 | 9496.006 | 0.603 | 0.437 |
| log(DNA in each size fraction/100) | 1 | 237 | 10088.420 | 593 | <0.001 |
| Size fraction | 2 | 93.2 | 9525.413 | 32.0 | <0.001 |
| Temperature | 1 | 85.2 | 9496.089 | 0.686 | 0.408 |
| Salinity | 1 | 85.7 | 9498.132 | 2.73 | 0.099 |
| Sampling location | 1 | 85.1 | 9495.723 | 0.320 | 0.572 |
| Total suspended solids | 1 | 85.6 | 9497.372 | 1.97 | 0.161 |
| Nitrate | 1 | 85.7 | 9498.039 | 2.64 | 0.104 |
| Silicate | 1 | 85.8 | 9498.274 | 2.87 | 0.090 |
| (Chlorophyll *a*/0.1) | 1 | 85.5 | 9497.015 | 1.61 | 0.204 |
| Size fraction:temperature | 2 | 88.1 | 9505.148 | 11.7 | 0.003 |
| Size fraction:salinity | 2 | 85.3 | 9494.317 | 0.914 | 0.633 |
| Size fraction:location | 2 | 89.5 | 9510.974 | 17.6 | <0.001 |
| Size fraction:total suspended solids | 2 | 93.7 | 9527.051 | 33.6 | <0.001 |
| Size fraction:nitrate | 2 | 85.1 | 9493.635 | 0.232 | 0.891 |

| | | | | | |
|---|---|---|---|---|---|
| Size fraction:silicate | 2 | 85.6 | 9495.615 | 2.21 | 0.331 |
| Size fraction:(chlorophyll *a*/0.1) | 2 | 92.7 | 9523.368 | 30.0 | <0.001 |
| **Within-seasons model: with monsoons included as a variable** | | | | | |
| Total Model | - | 70.7 | 9440.975 | - | - |
| DOC | 1 | 70.9 | 9440.015 | 1.04 | 0.308 |
| (Ammonia/0.1) | 1 | 71.4 | 9442.477 | 3.50 | 0.061 |
| (Nitrite/0.1) | 1 | 70.8 | 9439.418 | 0.443 | 0.506 |
| log(DNA in each size fraction/100) | 1 | 161 | 9898.859 | 460 | <0.001 |
| Size fraction | 2 | 83.2 | 9500.643 | 63.7 | <0.001 |
| Temperature | 1 | 71.5 | 9442.864 | 3.89 | 0.049 |
| Salinity | 1 | 70.9 | 9439.863 | 0.888 | 0.346 |
| Sampling location | 1 | 70.9 | 9439.865 | 0.889 | 0.346 |
| Monsoon seasons | 3 | 71.5 | 9439.043 | 4.07 | 0.254 |
| Total suspended solids | 1 | 71.4 | 9442.34 | 3.37 | 0.067 |
| Nitrate | 1 | 70.9 | 9439.988 | 1.01 | 0.314 |
| Silicate | 1 | 70.8 | 9439.674 | 0.699 | 0.403 |
| (Chlorophyll *a*/0.1) | 1 | 71.5 | 9443.227 | 4.25 | 0.039 |
| Size fraction:temperature | 2 | 72.5 | 9446.219 | 9.24 | 0.010 |
| Size fraction:salinity | 2 | 74.2 | 9454.523 | 17.5 | <0.001 |
| Size fraction:location | 2 | 73.5 | 9451.215 | 14.2 | <0.001 |
| Size fraction:monsoon | 6 | 79.2 | 9472.29 | 43.3 | <0.001 |
| Size fraction:total suspended solids | 2 | 75.3 | 9460.164 | 23.2 | <0.001 |
| Size fraction:nitrate | 2 | 73.0 | 9448.491 | 11.5 | 0.003 |
| Size fraction:silicate | 2 | 73.1 | 9449.187 | 12.2 | 0.002 |
| Size fraction:(chlorophyll *a*/0.1) | 2 | 72.8 | 9447.553 | 10.6 | 0.005 |

Table S4. Relative abundance and estimated abundance (*Vibrio* 16s rRNA gene copies/L) of the 20 most abundant *Vibrio* ASVs among the three size fractions at St. John's Island and Pulau Satumu. Estimated abundance was calculated based on the qPCR abundance of *Vibrio* spp. obtained in this study[1], and presented as the median abundance with the range indicated in parentheses. Where an ASV was undetected in any sample across the time series, it is indicated as below sequencing limit of detection (<LOD). The LOD was defined as at least one sequence detected within a sample.

| **St. John's Island** | | | | | | | |
|---|---|---|---|---|---|---|---|
| ASV | Top BLAST hit against NCBI Refseq database (% nucleotide identity) | **0.22–1 µm** | | **1–10 µm** | | **10–150 µm** | |
| | | % of *Vibrio* sequences | Estimated abundance from qPCR (range) | % of *Vibrio* sequences | Estimated abundance from qPCR (range) | % of *Vibrio* sequences | Estimated abundance from qPCR (range) |
| ASV_214 | *V. xuii* strain R-15052 (100%) | 7.13 | 11,361 (<LOD–242,130) | 9.68 | 27,194 (5341–493,840) | 14.8 | 6269 (415–82,155) |
| ASV_237 | *V. rotiferianus* CAIM 577=LMG 21460 (100%) | 8.20 | 13,010 (<LOD–246,624) | 10.5 | 41,438 (<LOD–406,973) | 12.9 | 6088 (452–96,107) |
| ASV_675 | *V. parahaemolyticus* strain ATCC 17802 (100%) | 6.09 | 10,783 (<LOD–116,096) | 7.06 | 17,915 (4644–150,188) | 6.49 | 2717 (200–20,435) |
| ASV_698 | *V. azureus* strain NBRC 104587 (99.5%) | 2.22 | 5083 (<LOD–53,470) | 3.85 | 18,481 (<LOD–77,384) | 4.18 | 1976 (279–13,928) |
| ASV_895 | *V. rumoiensis* strain S-1 (100%) | 0.323 | 672 (<LOD–33,187) | 2.50 | 12,768 (<LOD–54,088) | 4.38 | 3132 (212–13,811) |

[1] Good correspondence between qPCR abundance and *Vibrio* sequence relative abundance (see Fig. 5).

| | | | | | | | |
|---|---|---|---|---|---|---|---|
| ASV_897 | *V. caribbeanicus* ATCC BAA-2122 (100%) | 10.4 | 23,091 (6501–83,227) | 8.11 | 29,643 (5152–125,399) | 2.26 | 839 (36–10,525) |
| ASV_954 | *V. xiamenensis* strain G21 (99.26%) | 0 | 0 (<LOD –195,060) | 1.18 | 5465 (<LOD–259,397) | 1.43 | 737 (32–20,676) |
| ASV_975 | *V. brasiliensis* LMG 20546 (100%) | 2.02 | 6095 (<LOD –236,245) | 2.91 | 7735 (<LOD–312,196) | 0.522 | 75 (<LOD–20,901) |
| ASV_1114 | *V. japonicus* strain Bio7-2 (100%) | 1.55 | 3356 (<LOD–28,266) | 3.83 | 12,664 (2044–88,649) | 4.51 | 1257 (202–11,884) |
| ASV_1110 | *V. furnissii* strain 9119-82 (99.01%) | 0 | 0 (<LOD–118,066) | 0.797 | 1777 (<LOD–177,491) | 1.02 | 334 (<LOD–17,992) |
| ASV_1153 | *V. alginolyticus* strain NBRC 15630 (99.75%) | 0 | 0 (<LOD–9607) | 0 | 0 (<LOD–40,421) | 0 | 0 (<LOD–11,519) |
| ASV_1308 | *V. brasiliensis* LMG 20546 (99.5%) | 2.14 | 5479 (<LOD–77,669) | 2.18 | 10,918 (<LOD–123,406) | 1.70 | 671 (<LOD–10,094) |
| ASV_1340 | *V. caribbeanicus* ATCC BAA-2122 (99.75%) | 6.15 | 7306 (<LOD–43,480) | 0 | 0 (<LOD–78,439) | 0.225 | 128 (<LOD–5148) |
| ASV_1621 | *V. zhanjiangensis* strain E414 (100%) | 1.57 | 2665 (<LOD–42,507) | 1.57 | 4284 (<LOD–46,395) | 1.30 | 649 (59–5611) |
| ASV_1816 | *V. maritimus* strain R-40493 (99.75%) | 1.35 | 1876 (<LOD–33,460) | 1.92 | 5122 (<LOD–66,816) | 1.44 | 475 (<LOD–4263) |
| ASV_1902 | *V. rotiferianus* CAIM 577=LMG 21460 (99.75%) | 0 | 0 (<LOD–3799) | 0 | 0 (<LOD–26,844) | 0 | 0 (<LOD–4638) |

| | | | | | | | |
|---|---|---|---|---|---|---|---|
| ASV_2173 | *V. alginolyticus* strain NBRC 15630 (100%) | 2.12 | 3770 (<LOD–18,498) | 0.833 | 2421 (<LOD–32,583) | 1.28 | 140 (<LOD–3884) |
| ASV_1786 | *V. mediterranei* strain 50 (100%) | 2.22 | 4441 (<LOD–39,128) | 1.76 | 7399 (<LOD–40,813) | 0.288 | 271 (<LOD–1668) |
| ASV_2110 | *V. rotiferianus* CAIM 577=LMG 21460 (99.51%) | 0 | 0 (<LOD–41,109) | 0 | 0 (<LOD–35,934) | 0.729 | 180 (<LOD–10,752) |
| ASV_2209 | *V. zhanjiangensis* strain E414 (99.75%) | 0.233 | 495 (<LOD–26,883) | 1.11 | 5527 (<LOD–25,308) | 0.938 | 391 (<LOD–3162) |
| **Pulau Satumu** | | | | | | | |
| ASV | Top BLAST hit against NCBI Refseq database (% nucleotide identity) | **0.22–1 µm** | | **1–10 µm** | | **10–150 µm** | |
| | | % of *Vibrio* sequences | Estimated abundance from qPCR (range) | % of *Vibrio* sequences | Estimated abundance from qPCR (range) | % of *Vibrio* sequences | Estimated abundance from qPCR (range) |
| ASV_214 | *V. xuii* strain R-15052 (100%) | 4.14 | 3763 (<LOD–762,591) | 7.45 | 20,741 (2507–990,139) | 13.1 | 1493 (357–22,265) |
| ASV_237 | *V. rotiferianus* CAIM 577=LMG 21460 (100%) | 8.31 | 24,677 (<LOD–544,061) | 14.2 | 25,297 (3885–713,551) | 18.1 | 1711 (274–16,351) |
| ASV_675 | *V. parahaemolyticus* strain ATCC 17802 (100%) | 4.15 | 9630 (<LOD–102,477) | 4.75 | 16,391 (648–131,196) | 7.07 | 917 (176–15,685) |
| ASV_698 | *V. azureus* strain NBRC 104587 (99.5%) | 2.17 | 3928 (<LOD–39,901) | 4.22 | 11,988 (<LOD–83,013) | 9.41 | 594 (272–14,889) |
| ASV_895 | *V. rumoiensis* strain S-1 (100%) | 0.602 | 487 (<LOD–35,684) | 1.15 | 3968 (830–40,918) | 2.76 | 181 (<LOD–5212) |

| | | | | | | | |
|---|---|---|---|---|---|---|---|
| ASV_897 | *V. caribbeanicus* ATCC BAA-2122 (100%) | 13.4 | 26,457 (11,946–146,739) | 8.09 | 24,622 (6886–146,876) | 2.86 | 333 (9–6990) |
| ASV_954 | *V. xiamenensis* strain G21 (99.26%) | 0.541 | 802 (<LOD–100,044) | 0.861 | 2316 (<LOD–50,056) | 0.910 | 89 (<LOD–886) |
| ASV_975 | *V. brasiliensis* LMG 20546 (100%) | 0 | 0 (<LOD–85,172) | 0.639 | 2052 (<LOD–87,009) | 0.223 | 47 (<LOD–821) |
| ASV_1114 | *V. japonicus* strain Bio7-2 (100%) | 1.35 | 1147 (<LOD–35,968) | 2.39 | 6140 (415–74,423) | 3.18 | 441 (57–2695) |
| ASV_1110 | *V. furnissii* strain 9119-82 (99.01%) | 0.377 | 988 (<LOD–41,116) | 0.619 | 1317 (<LOD–46,859) | 0.392 | 84 (<LOD–1066) |
| ASV_1153 | *V. alginolyticus* strain NBRC 15630 (99.75%) | 0 | 0 (<LOD–10,575) | 0 | 0 (<LOD–24,644) | 0 | 0 (<LOD–5549) |
| ASV_1308 | *V. brasiliensis* LMG 20546 (99.5%) | 0 | 0 (<LOD–50,865) | 0.909 | 1089 (<LOD–50,146) | 0.399 | 104 (<LOD–332) |
| ASV_1340 | *V. caribbeanicus* ATCC BAA-2122 (99.75%) | 4.65 | 9723 (<LOD–100,463) | 2.98 | 11,008 (<LOD–51,769) | 0.750 | 97 (<LOD–1341) |
| ASV_1621 | *V. zhanjiangensis* strain E414 (100%) | 0.680 | 1309 (<LOD–23,183) | 1.64 | 3034 (<LOD–24,217) | 0.802 | 135 (<LOD–667) |
| ASV_1816 | *V. maritimus* strain R-40493 (99.75%) | 1.87 | 3936 (<LOD–78,843) | 1.71 | 4865 (923–93,678) | 1.39 | 110 (<LOD–420) |
| ASV_1902 | *V. rotiferianus* CAIM 577=LMG 21460 (99.75%) | 0 | 0 (<LOD–13,306) | 0 | 0 (<LOD–18,314) | 0 | 0 (<LOD–3812) |

| | | | | | | | |
|---|---|---|---|---|---|---|---|
| ASV_2173 | *V. alginolyticus* strain NBRC 15630 (100%) | 2.04 | 3763 (<LOD–13,268) | 1.45 | 5096 (<LOD–22,235) | 1.27 | 133 (<LOD–21,846) |
| ASV_1786 | *V. mediterranei* strain 50 (100%) | 2.74 | 3186 (<LOD–10,7428) | 1.07 | 2555 (<LOD–102,349) | 0.625 | 198 (<LOD–622) |
| ASV_2110 | *V. rotiferianus* CAIM 577=LMG 21460 (99.51%) | 0 | 0 (<LOD–17,114) | 0 | 0 (<LOD–31,786) | 0.320 | 35 (<LOD–497) |
| ASV_2209 | *V. zhanjiangensis* strain E414 (99.75%) | 0.541 | 988 (<LOD–16,874) | 0.824 | 3227 (<LOD–25,704) | 0.712 | 112 (19–592) |

Table S5. Imputation-related variance in the pooled comparison of across-seasons and within-seasons models. The proportion of total test variance attributable to imputation uncertainty (lambda) are shown for the four abundant *Vibrio* ASVs most closely related to human pathogens. Below-LOD *Vibrio* values were imputed across 15 completed datasets; larger lambda values indicate greater dependence of the pooled comparison on the imputation step.

| ASV | lambda |
|---|---|
| *Vpa_675* | 0.029 |
| *Vfu_1110* | 0.238 |
| *Val_2173* | 0.229 |
| *Val_1153* | 0.392 |

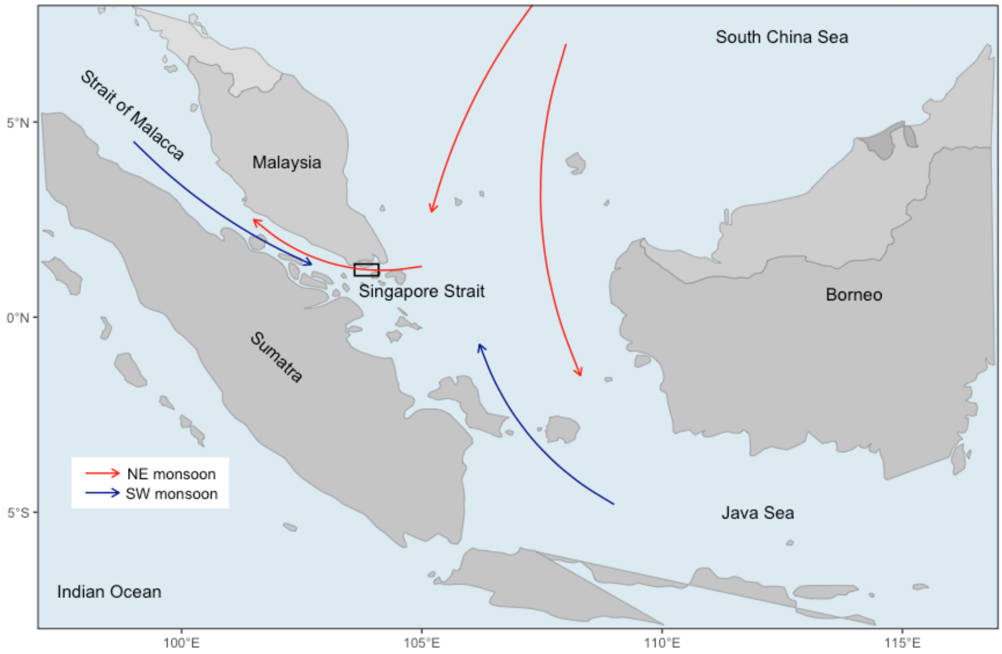


Fig. S1. Map of Singapore and neighboring countries and territories showing the seasonal reversal of ocean currents between South China Sea and Strait of Malacca through the Singapore Strait during the two monsoon periods. The Singapore Strait is demarcated by the black box. Adapted from Poo et al. (2026).

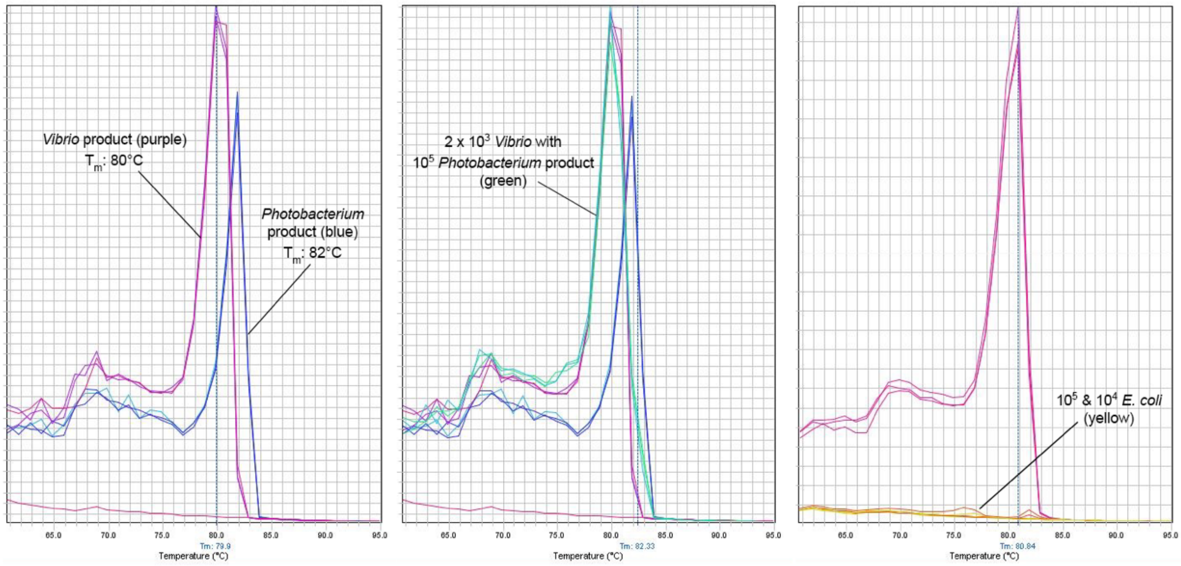


Fig. S2. Melt curve profiles of the 567F/680R qPCR assay amplifying the corresponding 16s rRNA target fragment from *Vibrio*, *Photobacterium* (closely related genus to *Vibrio*) and *E. coli*. Annealing temperature at 70ºC achieved assay specificity to *Vibrio* sequences. The melt curves were generated by the StepOnePlus™ Software v2.3.

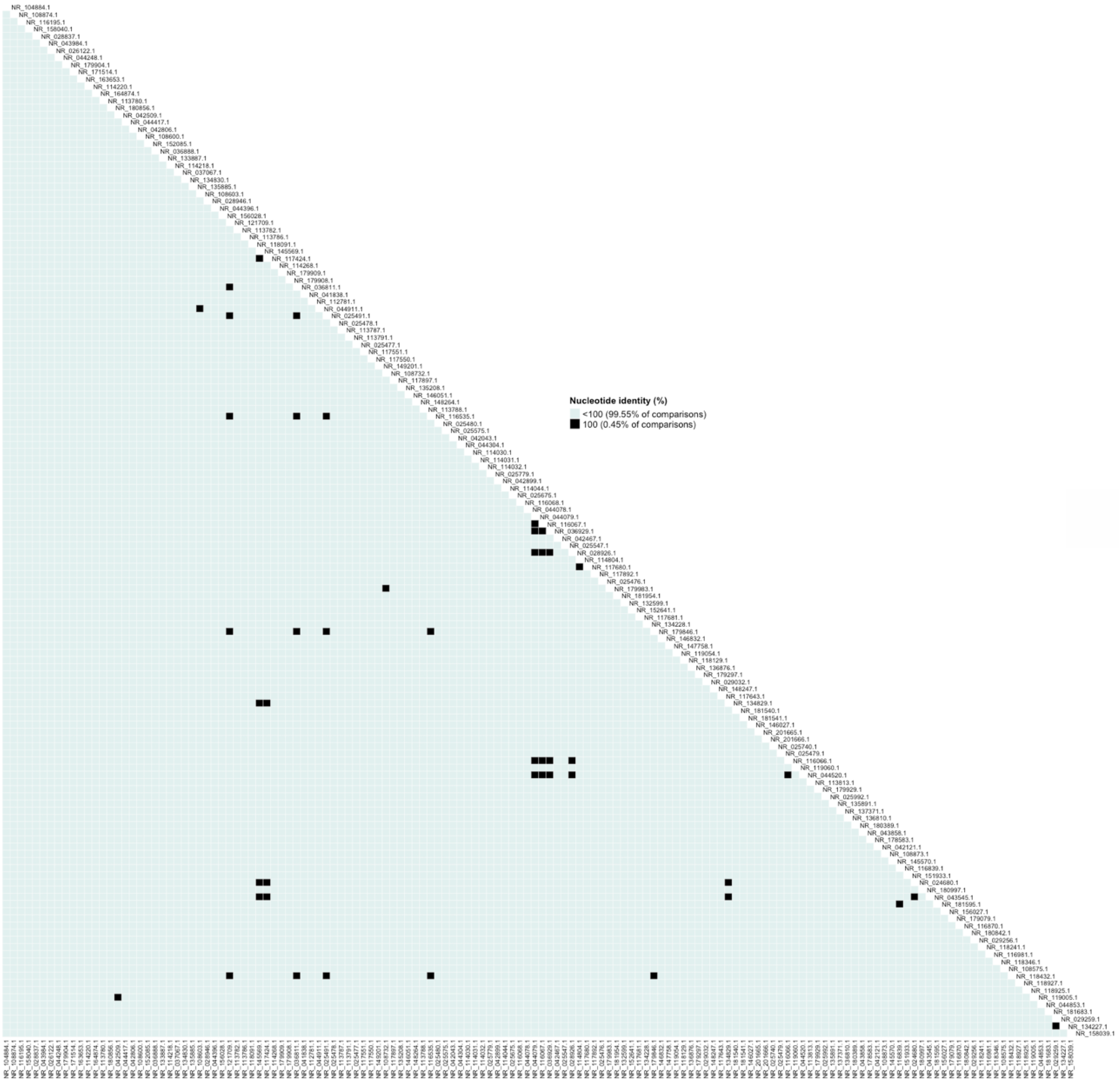


Fig. S3. Pairwise comparisons of nucleotide identity over the amplified 404–405 nt V6-8 region among 144 *Vibrio* sequences from the NCBI Reference Sequence database, showing good performance of the V6-8 region in distinguishing species-level diversity in *Vibrio*[2]. The 144 reference sequences are represented by their accession numbers. Pairs with 100% nucleotide identity of this amplified region are indicated in black (46 pairs, 0.45%) while pairs with <100% identity (i.e. at least one nucleotide difference) are depicted in green (10 250 pairs, 99.55%). Self-comparisons are not shown for figure clarity.

---

[2] Acknowledging that some species have multiple sequences in this region due to intra-genomic operon diversity

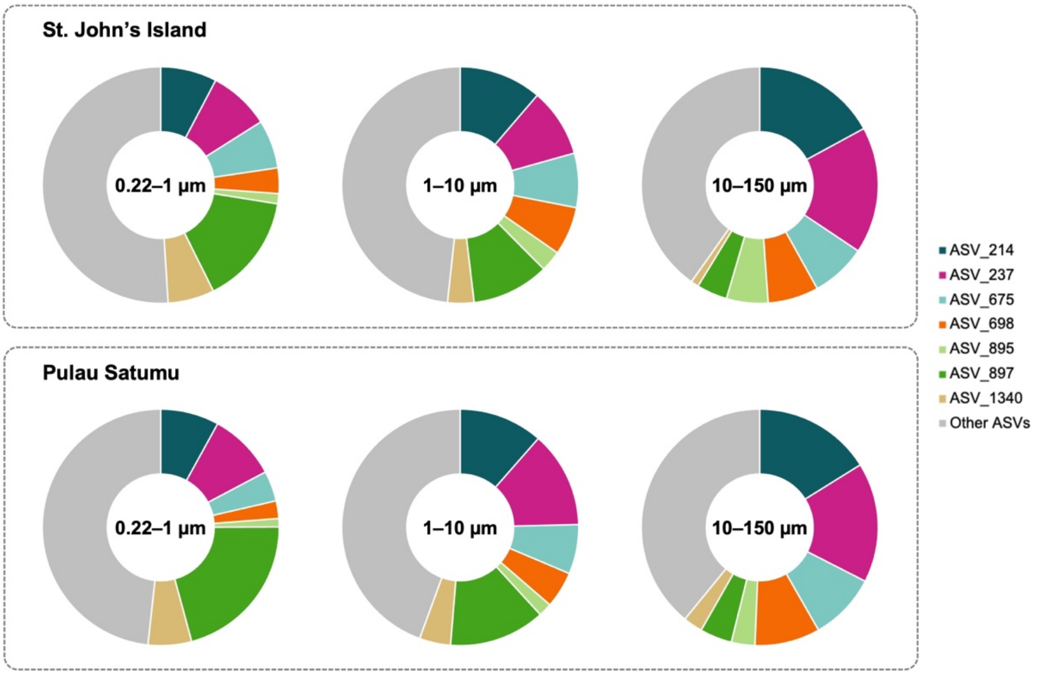


Fig. S4. Composition of the *Vibrio* assemblage (by relative abundance) among the three size fractions at St. John's Island and Pulau Satumu. The pie charts indicate average proportion of ASVs (relative to all sequences belonging to *Vibrio*) over the time series.

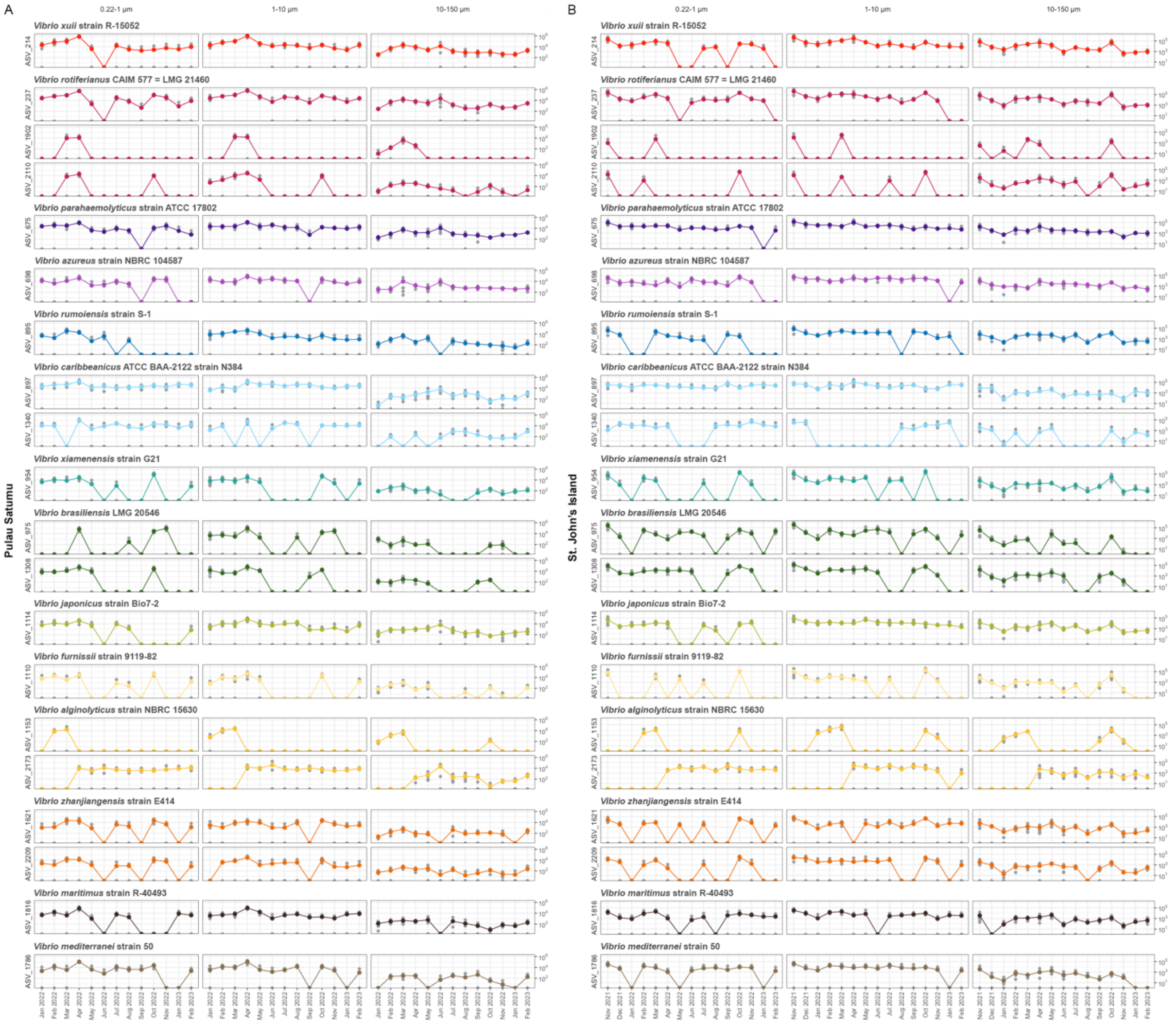


Fig. S5. Abundance estimates of the 20 most abundant *Vibrio* ASVs among the three size fractions at (**A**) Pulau Satumu and (**B**) St. John's Island. Abundance was calculated based on the relative abundances of individual ASVs from the time series 16s rRNA amplicon dataset and the qPCR abundance of *Vibrio* spp. obtained in this study, and presented as the mean (with $N$=5 per time point and sampling location indicated with grey data points). Each ASV is labelled with its top BLAST hit (against the Refseq database) and similarly annotated ASVs are grouped together.

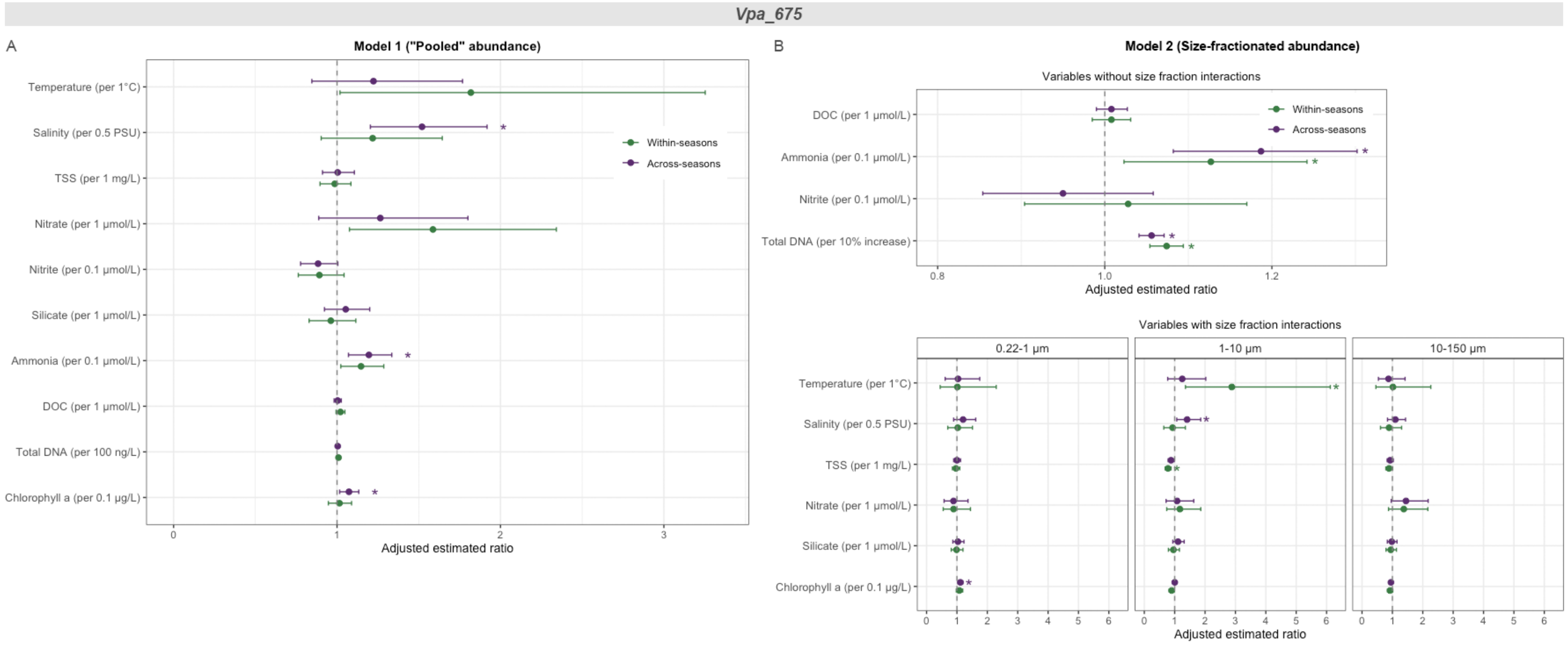


Fig. S6. Predictor effects of environmental variables from GLM analysis of *Vpa_675*. Predictor effects were presented as adjusted estimated ratio with 95% confidence intervals for (**A**) "pooled" abundance across all size fractions (Model 1) and (**B**) abundance in each of the three size fractions (Model 2) and compared between the across-seasons (without accounting for monsoons) and within-seasons (that included 'monsoon seasons' as a variable) model. Each trend is estimated while holding all other variables constant. All comparisons have been adjusted for multiple comparisons using the Benjamini-Hochberg method. Variables that were significant predictors ($p$<0.05) in each model are indicated with asterisks.

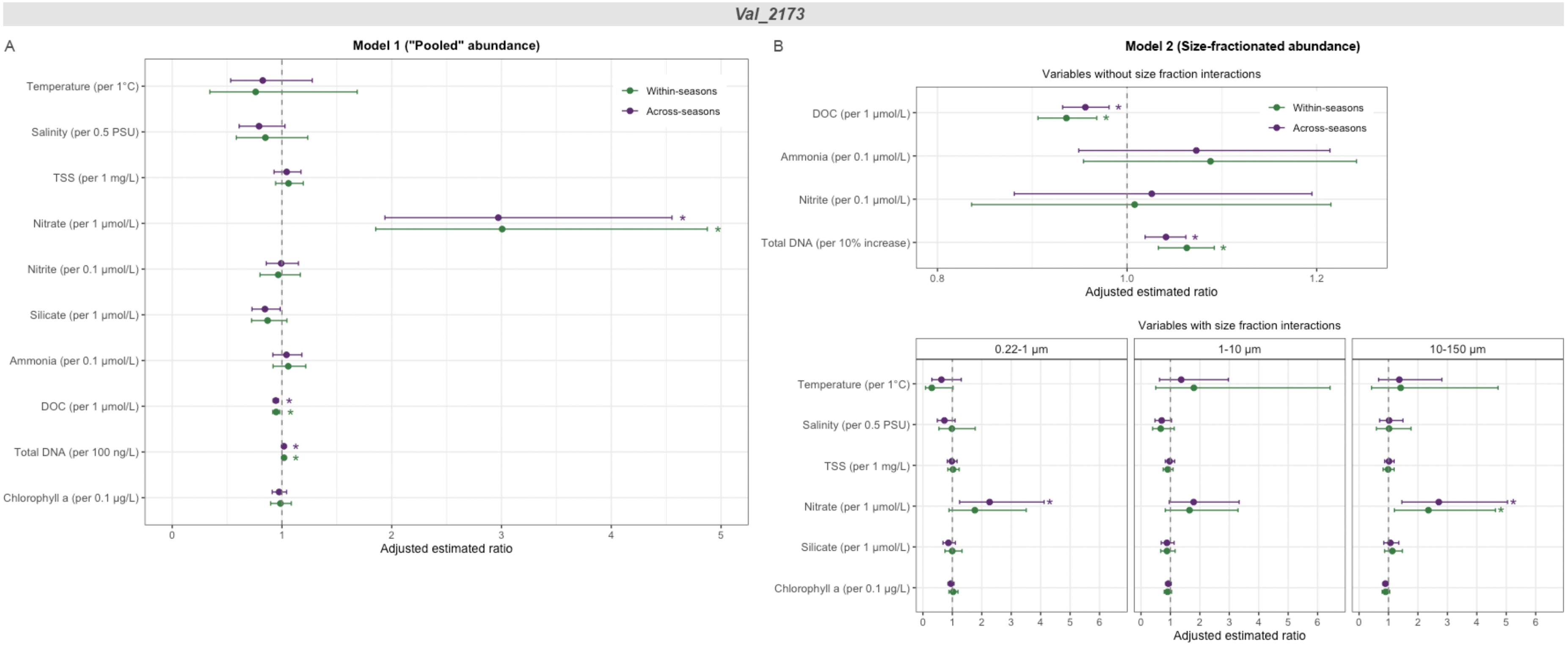


Fig. S7. Predictor effects of environmental variables from GLM analysis of *Val_2173*. Predictor effects were presented as adjusted estimated ratio with 95% confidence intervals for (**A**) "pooled" abundance across all size fractions (Model 1) and (**B**) abundance in each of the three size fractions (Model 2) and compared between the across-seasons (without accounting for monsoons) and within-seasons (that included 'monsoon seasons' as a variable) model. Each trend is estimated while holding all other variables constant. All comparisons have been adjusted for multiple comparisons using the Benjamini-Hochberg method. Variables that were significant predictors ($p$<0.05) in each model are indicated with asterisks.

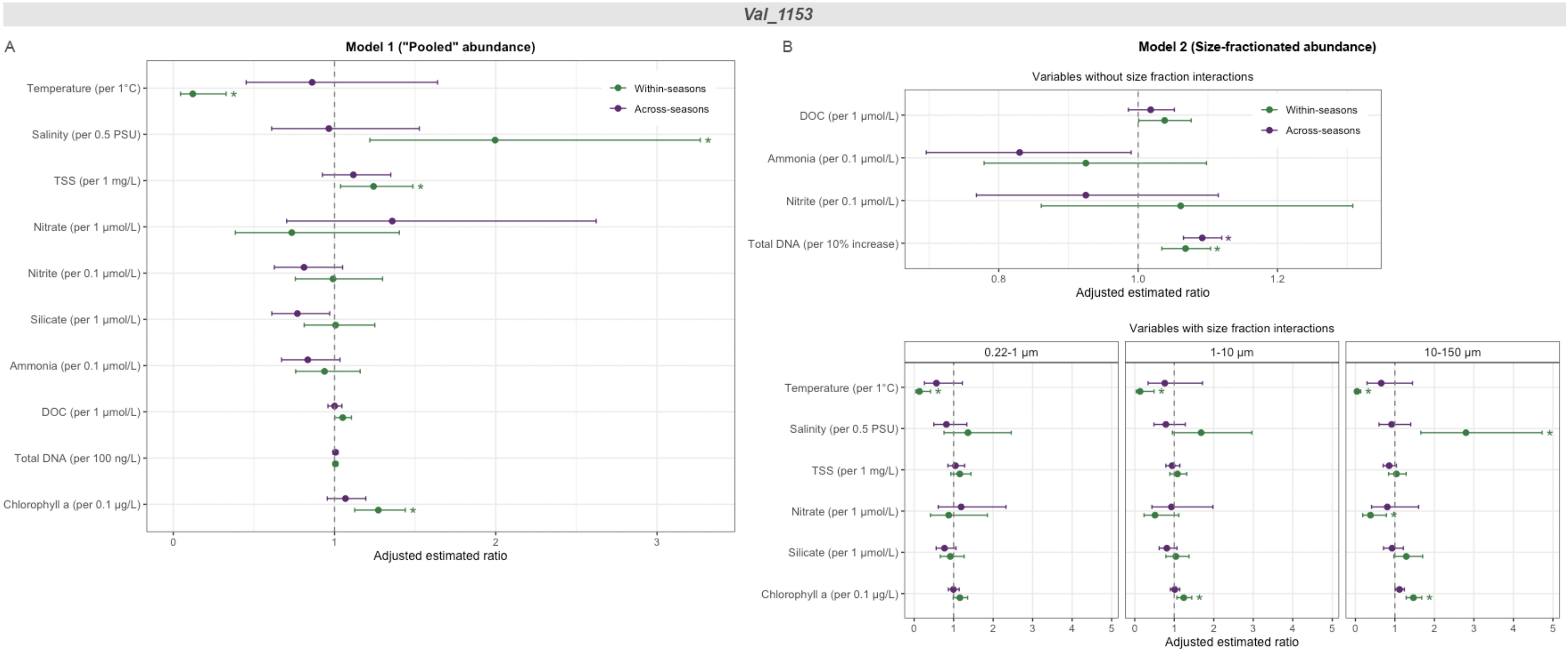


Fig. S8. Predictor effects of environmental variables from GLM analysis of *Val_1153*. Predictor effects were presented as adjusted estimated ratio with 95% confidence intervals for (**A**) "pooled" abundance across all size fractions (Model 1) and (**B**) abundance in each of the three size fractions (Model 2) and compared between the across-seasons (without accounting for monsoons) and within-seasons (that included 'monsoon seasons' as a variable) model. Each trend is estimated while holding all other variables constant. All comparisons have been adjusted for multiple comparisons using the Benjamini-Hochberg method. Variables that were significant predictors ($p$<0.05) in each model are indicated with asterisks.

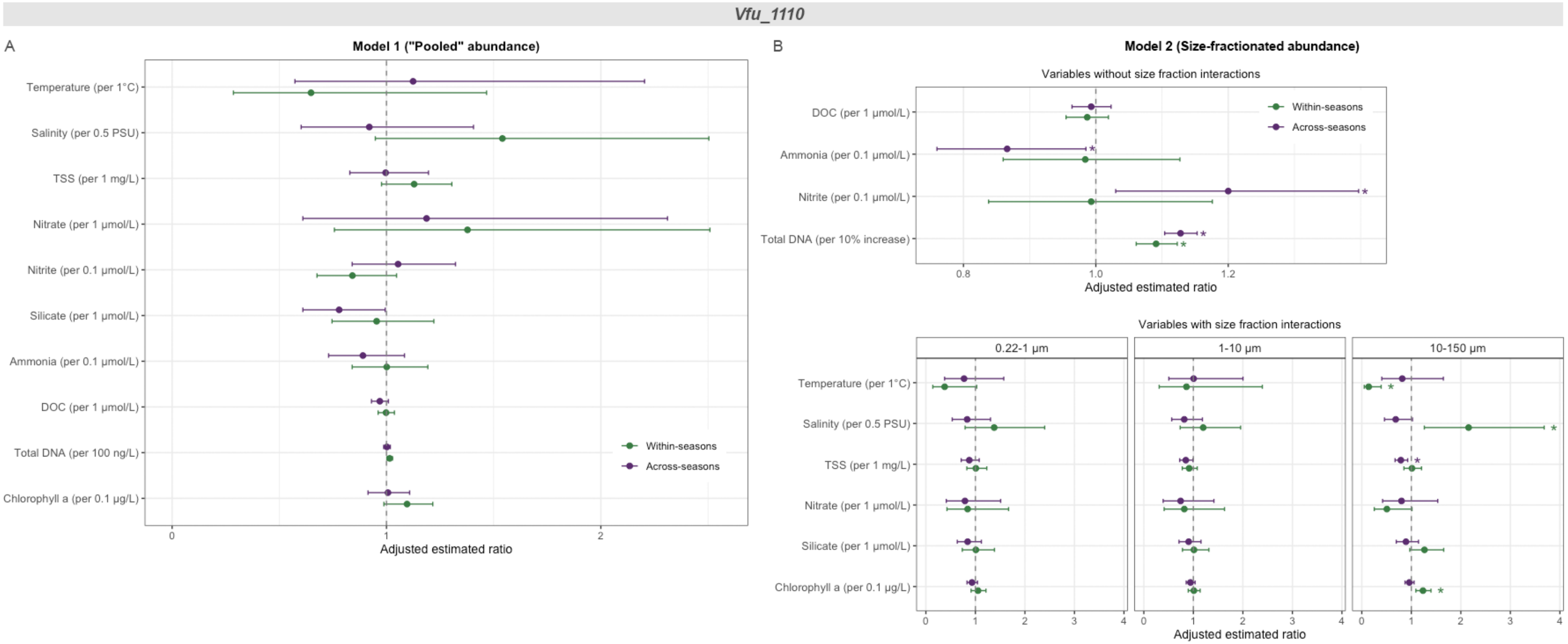


Fig. S9. Predictor effects of environmental variables from GLM analysis of *Vfu_1110*. Predictor effects were presented as adjusted estimated ratio with 95% confidence intervals for (**A**) "pooled" abundance across all size fractions (Model 1) and (**B**) abundance in each of the three size fractions (Model 2) and compared between the across-seasons (without accounting for monsoons) and within-seasons (that included 'monsoon seasons' as a variable) model. Each trend is estimated while holding all other variables constant. All comparisons have been adjusted for multiple comparisons using the Benjamini-Hochberg method. Variables that were significant predictors ($p$<0.05) in each model are indicated with asterisks.
